# Chirality-Enhanced Transport and Drug Delivery of Graphene Nanocarriers to Tumor-like Cellular Spheroid

**DOI:** 10.1101/2023.06.27.546698

**Authors:** Hyunsu Jeon, Runyao Zhu, Gaeun Kim, Yichun Wang

## Abstract

Chirality, defined as “a mirror image,” is a universal geometry of biological and nonbiological forms of matter. This geometry of molecules determines how they interact during their assembly and transport. With the development of nanotechnology, many nanoparticles with chiral geometry or chiroptical activity have emerged for biomedical research. The mechanisms by which chirality originates and the corresponding synthesis methods have been discussed and developed in the past decade. Inspired by the chiral selectivity in life, a comprehensive and in-depth study of interactions between chiral nanomaterials and biological systems has far-reaching significance in biomedicine. Here, we investigated the effect of the chirality of nanoscale drug carriers, graphene quantum dots (GQDs), on their transport in tumor-like cellular spheroids. Chirality of GQDs (*L/D*-GQDs) was achieved by the surface modification of GQDs with *L/D*-cysteines. As an *in-vitro* tissue model for drug testing, cellular spheroids were derived from a human hepatoma cell line (*i*.*e*., HepG2 cells) using the Hanging-drop method. Our results reveal that the *L*-GQDs had a 1.7-fold higher apparent diffusion coefficient than the *D*-GQDs, indicating that the left-handed chirality of GQDs can enhance their transport into tumor-like cellular spheroids. Moreover, when loaded with a common chemotherapy drug, Doxorubicin (DOX), *via* π-π stacking, *L*-GQDs are more effective as nanocarriers for drug delivery, resulting in 25% higher efficacy for cancerous cellular spheroids than free DOX. Overall, our studies indicated that the chirality of nanocarriers is essential for the design of drug delivery vehicles to enhance the transport of drugs in a cancerous tumor.

## 1. Introduction

The left- or right-handedness of molecules (*e*.*g*., *L*- or *D*-), known as chirality, plays a crucial role in biological processes (Zhao et al., 2020; Shao et al., 2021). The interaction of biomolecules depends on their three-dimensional (3D) shape and spatial arrangement, while typically, one type of chiral molecules (*i*.*e*., enantiomers) is biologically active, constructing biopolymers as homochiral building blocks (Salam, 1991; Warning et al., 2021). This rule applies not only to natural molecules in living organisms (*e*.*g*., amino acids, nucleotides, sugars, their oligomers, and macromolecules (Powell et al., 2012)) but also to synthetic matters, including both small chemicals (*i*.*e*., drugs (Speirs, 1962; Du et al., 2020; Yeom et al., 2020)) and bulk substances (*i*.*e*., biomaterials (Wang et al., 2019)). With the development of nanotechnology, many nanoparticles (NPs) with chiral geometry or chiroptical activity have emerged for biomedical applications (Suzuki et al., 2016; Du et al., 2017; Jiang et al., 2017; Yeom et al., 2020; Peng et al., 2021; Döring et al., 2022; Wang et al., 2022b; Wu and Pauly, 2022). The origin of chirality and the associated synthesis techniques have been extensively explored and refined in previous studies (Suzuki et al., 2016; Jiang et al., 2017; Ma et al., 2017; Döring et al., 2022; Wu and Pauly, 2022). Drawing inspiration from the selectivity exhibited by chiral entities in biological systems, a thorough and rigorous investigation into the nanoscale interactions between chiral nanomaterials and biological systems have profound implications for biomedicine. Recent studies showed that the chiral interaction of NPs with protein and lipid membranes at bio-interfaces plays a key role in cell uptake (Shanker and Aschner, 2001; Shanker et al., 2001; Al-Hajaj et al., 2011) and tissue transport (Du et al., 2017; Huang et al., 2020a). For instance, phospholipid bilayers, the fundamental structure of the cell membrane, are enantio-selectively permeable to chiral nanomaterials due to their chiral interaction, which results in highly efficient cell uptake of right-handed (*D-*) NPs with nanocore-dominant chirality (Sarasij et al., 2007; Suzuki et al., 2016; Du et al., 2017; Yeom et al., 2020). Meanwhile, chiral NPs interacting with membrane proteins exhibit asymmetric active transport through cell membranes (Sarasij et al., 2007). Left-handed (*L*-) NPs with ligand-dominant chirality affect active cellular uptake in human cells as well as cellular efflux (Al-Hajaj et al., 2011). Lastly, the transport and distribution of NPs in tissues are strongly influenced by the dynamics of NPs interacting with the cell membrane and extracellular matrix (ECM) (Sherman and Fisher, 1986; Cooper, 2000; Ng and Pun, 2008; Chauhan et al., 2009; Hsu et al., 2012; Gao et al., 2013; Wang et al., 2015; Ziemys et al., 2018; Fulaz et al., 2019; Leedale et al., 2020; Lenzini et al., 2020; Koomullil et al., 2021). Engineering chirality of NPs can enhance their transport and uptake in the tissue microenvironment by altering the reaction-diffusion processes due to the nanocarrier-induced cell uptake and non-diffusive active transport due to the proteinaceous polyelectrolyte disassembly on the tissue ECM (Huang et al., 2020b). Hence, it is essential to consider the chirality of nanocarriers and understand their effect on the design of drug delivery systems (Zhao et al., 2021).

Graphene quantum dots (GQDs), a graphene-based NP, have emerged as a versatile and promising tool for bio-medical applications in drug delivery (Yeom et al., 2020), biosensing (Hai et al., 2018), and bioimaging (Li et al., 2023), due to their unique properties, including high surface area-to-volume ratio (Tian et al., 2018), tunable optical properties (Hai et al., 2018), and excellent biocompatibility (Henna and Pramod, 2020). For example, GQDs emit strong, stable, and tunable fluorescence signals (Wang et al., 2016) for high-resolution imaging of biological structures (Biswas et al., 2021). They can be easily functionalized with specific biomolecules to enhance their biostability (Vázquez-Nakagawa et al., 2022), endow desired functionality (Suzuki et al., 2016; Sattari et al., 2021), target specific cells or tissues (Yao et al., 2017), or contain chemical drugs (Sawy et al., 2021). Notably, the planar plane of GQDs mainly consists of aromatic rings, which makes them well-suited for loading small aromatic molecules, such as chemotherapy drugs (Askari et al., 2021; Mirzaei-Kalar et al., 2022), through aromatic stacking (*i*.*e*., π-π stacking) (Liu et al., 2009). This property enabled the development of GQDs as drug carriers for drug delivery that could effectively transport and release the loaded drugs to targeted cells or tissues with high selectivity and efficacy (Liu et al., 2009; Wo et al., 2016; Askari et al., 2021; Kiani Nejad et al., 2022; Mirzaei-Kalar et al., 2022). The edges of GQDs can be engineered to contain desired functional groups, which enables covalent modification with chiral molecules, endowing nanoscale chirality (Suzuki et al., 2016). Moreover, chiral matching with the cellular lipid membranes (Sarasij et al., 2007; Suzuki et al., 2016; Du et al., 2017; Yeom et al., 2020) enhanced the loading of chiral GQDs with embedded chemicals into extracellular vesicles for achieving advanced exosome engineering (Zhang et al., 2023).

This study investigated how the chirality of GQDs affected their transport in tumor-like cellular spheroids, a well-known 3D cell culture model (**Figure 1**). Compared to the traditional two-dimensional (2D) cell culture models, 3D cell culture mimics the *in vivo* tissue microenvironment morphologically and compositionally (Wang and Jeon, 2022) by replicating *in vivo* cell growth, proliferation, and differentiation *in vitro* (Charoen et al., 2014; Chen, 2016), providing insight into nanocarrier transport in native tissues (Gunti et al., 2021). The chirality of GQDs was obtained by surface modification with *L/D*-cysteines (**Figure 1A**), which gave a strong chiroptical activity of GQDs and indicated nanoscale chirality by the twist of the graphene nanosheets. (Wang et al., 2016; Zhang et al., 2023). The chiral GQDs were tested in the tumor-like cellular spheroids, which were derived from a human hepatoma cell line (HepG2) by Hanging-drop (*i*.*e*., 3-day-cultured cellular aggregates) (Timmins and Nielsen, 2007) followed by suspension culture (*i*.*e*., 10-day-cultured cellular spheroids) (Ryu et al., 2019) (**Figure 1B**). From the results, we found chirality-dependent phenomena in both cellular aggregates and spheroids: **1)** while tested in cellular aggregates, *L/D*-GQDs interacted with the immature ECM of cellular aggregates differently, occurring distinct structural change (*i*.*e*., swelling) of cellular aggregates (**Figure 1C**); **2)** we observed faster transport of *L*-GQDs into cellular spheroids compared to that of *D*-GQDs, representing significant differences in apparent diffusion coefficients in tumor-like tissue (**Figure 1D**). Accordingly, as a proof of concept for drug delivery application, DOX-loaded *L*-GQDs showed increased efficacy in tumor-like cellular spheroids compared to free DOX or DOX-loaded *D*-GQDs, conducting enhanced drug transport and delivery to the cancerous tissue model (**Figure 1E**). In summary, the chirality of GQDs influences their transport into tumor-like tissues, thus their delivery efficiency as nanocarriers, which indicates the importance and potential of chirality for designing and developing nanocarriers for drug delivery systems.

**Figure 1.**
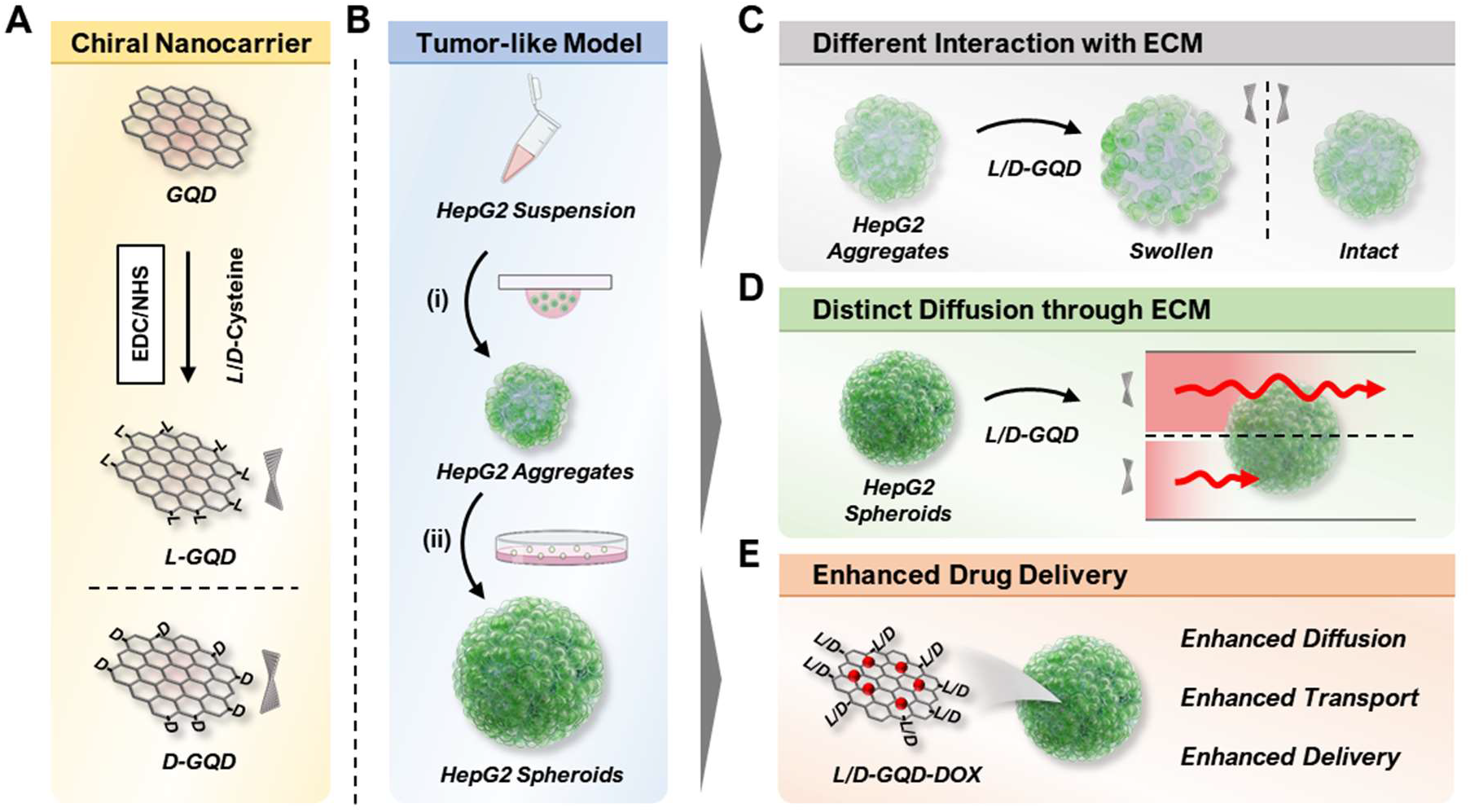
Illustration of overall workflow. **(A)** Scheme of the left- or right-handed graphene quantum dots (*L/D*-GQDs) preparation. **(B)** Scheme showing the human hepatoma (HepG2) cellular spheroid preparation. The cellular aggregates were prepared by **(i)** the Hanging drop method and matured to cellular spheroids by **(ii)** the suspension culture method. **(C)** Graphene quantum dots (GQDs) with chirality showed a distinct effect on the structure change of cellular aggregates (*e*.*g*., swelling). **(D)** Chiral GQDs showed distinct diffusivities in HepG2 cellular spheroids. **(E)** Chiral GQDs can serve as a nanocarrier for doxorubicin (DOX) with enhanced diffusion, transport, and delivery in tumor-like tissue.

## 2. Materials and Methods

### 2.1 Materials and Agents

Carbon nanofibers (719803) were purchased from Sigma-Aldrich (MO), and sulfuric acid (95-98%; BDH3068-500MLP) and nitric acid (69-70%; BDH3044-500MLPC) were purchased from VWR (PA). The dialysis membrane tubing (MWCO: 1 kD; 20060186) was purchased from Spectrum Chemical Manufacturing Company (NJ). 1-ethyl-3-(3-dimethyl-aminopropyl) carbodiimide (EDC; 22980) was purchased from Thermofisher Scientific (MA). N-hydroxy-succinimide sodium salt (Sulfo-NHS; 56485) and *L*-cysteine (30089) were purchased from Sigma-Aldrich (MO). *D*-cysteine (A110205-011) was purchased from AmBeed (IL).

Minimum essential medium Eagle 1 X (MEM; 10-010-CV) and phosphate-buffered saline 1 X (PBS; 21-040-CM) were purchased from Corning (NY). Avantor Seradigm USDA-approved origin fetal bovine serum (FBS; 1300-500) was purchased from VWR (PA), kept at temperature of -20°C (*e*.*g*., without the heat inactivation). Antibiotic antimycotic (15240096) was purchased from Fisher Scientific (MA). Gibco™ Trypsin-EDTA (25200072) was purchased from Thermofisher Scientific (MA). 4% Paraformaldehyde in 0.1 M phosphate buffer (15735-50S) was purchased from Electron Microscopy Sciences (PA). Ethanol absolute (64-17-5) was purchased from VWR (PA). LIVE/DEAD™ Cell Imaging Kit (488/570) (R37601) was purchased from Thermofisher Scientific (MA).

### 2.2 GQD Synthesis and Functionalization

GQDs were prepared using a modified Hummers method (Peng et al., 2012). 0.45 g of carbon fiber was added into 90 mL of concentrated H_2_SO_4_ (98%) and stirred for 1.5 h in a 250 mL round bottom glass flask. After stirring, 30 mL of concentrated HNO_3_ (68%) was added to the mixture solution and sonicated for 1 h in a 250 mL round bottom glass flask. Then, the mixture reacted at 120°C for 20 h. Next, the solution was neutralized by a 5 M of sodium hydroxide solution. The final product was further dialyzed for 3 days in a dialysis bag with 1K MWCO for purification.

The surface modification of GQDs with cysteines (*e*.*g*., *L*-GQDs and *D*-GQDs) was carried out using EDC/NHS coupling reaction (Suzuki et al., 2016; Park and Park, 2022). 1 mL of EDC with a concentration of 100 mM (in DI water) was mixed with 25 mL GQDs (0.25 mg·mL^−1^ in DI water), Followed by 30-min stirring. 1 mL of NHS (500 mM in DI water) was added into the mixture solution and stirred for 30 min. 1 mL of *L/D*-cysteines (100 mM in DI water) was added into the reaction, and the mixture reacted for 16 h. The product was purified by a dialysis bag with 1 K MWCO followed by filtration with VWR syringe filter (0.22 μm; 76479-010; VWR, PA). To load DOX to GQDs (*e*.*g*., *L*-GQD-DOX or *D*-GQD-DOX), 0.5 mg·mL^-1^ of GQDs and 350 μM of DOX were prepared under a deionized water solution followed by room-temperature incubation for 30 min protected from light. The DOX loading efficiency test was performed by measuring the fluorescence intensity of 200 μM DOX loaded in a concentration range of *L/D*-GQDs (*e*.*g*., 0, 75, 150, 225, and 300 μg·mL^-1^, respectively). The measurement was performed with Tecan Infinite 200Pro (Tecan, Männedorf, Switzerland) under 480 nm excitation for 600 nm emission. The loading efficiency was calculated by the following equation (1), where η is the loading efficiency, *I*_*0*_ is fluorescence intensity from unquenched DOX, *I*_*Q*_ is fluorescence intensity from quenched DOX, *C* is concentration of *L/D*-GQDs, and *I(C)* is DOX fluorescence intensity as a function of GQD concentration (*e*.*g*., *C*).

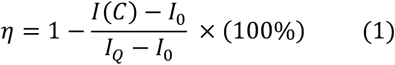

### 2.3 GQD Characterization

The size and shape of *L/D*-GQDs were characterized by a transmission electron microscope (TEM; JEOL 2011; Joel, WA). A 3-μL droplet of the *L/D*-GQD solution (0.1 mg·mL^-1^) was placed on the carbon-coated copper TEM grid (CF200-Cu-25; Purchased from Electron Microscopy Sciences, PA) and allowed to dry in the air. The imaging was performed with a TEM instrument under an accelerating voltage of 200 kV. The size distribution of the GQDs was analyzed by using ImageJ software. The chemical composition of *L/D*-GQDs was estimated with Fourier transform infrared (FTIR) spectroscopy, compared to the unmodified GQDs. Attenuated Total Reflectance (ATR)-FTIR analysis was performed using a Bruker Tensor 27 FTIR Spectrometer (Bruker Optics International Company, MA) with a diamond lens ATR module. 3 μL of 2 mg·mL^−1^ GQDs was dried in the air, and each spectrum was measured as the accumulation of 64 scans at a spectral resolution of 2 cm^−1^ within the range 4000–700 cm^−1^.

The absorbance and circular dichroism (CD) spectra of *L/D*-GQDs were evaluated with CD spectroscopy (Jasco J-1700 Spectrometer; Jasco International Company, MD) at 20°C. The *L*/*D*-GQD solution was diluted to 50 μg·mL^-1^, followed by spectra scanning from 200 nm to 400 nm with 0.1 nm intervals, 5 nm bandwidth, and a scan speed of 50 nm·min^−1^. The fluorescence emission property of *L/D*-GQDs was measured by Tecan Infinite 200Pro (Tecan, Männedorf, Switzerland). The fluorescence profile of the *L/D*-GQD solution was measured throughout a range of emission wavelengths from 400 nm to 700 nm under the 365 nm excitation.

### 2.4 Cell Culture and Cellular Spheroid Fabrication

Hepatocellular carcinoma human cells (HepG2) (American Type Culture Collection (ATCC), VA) were maintained with MEM supplemented with 10% FBS and 1% Antibiotic-Antimyotic in a humidified incubator (MCO-15AC; Sanyo, Osaka, Japan) at 37°C with 5% CO_2_. The cells were passaged routinely to maintain exponential growth (*e*.*g*., Passages 6-12). HepG2 cells were washed with PBS, trypsinized with trypsin-EDTA solution, and condensed at 3.4 × 10^4^ cells·mL^−1^ in a fresh cell culture medium. 7-μL droplets of cell suspension were gently arrayed on the lid of a 60 mm non-treated culture dish. The bottom of the dish was filled with 6 mL of PBS solution, followed by the droplet-arrayed lid covering. The culture dish was incubated for 72 h at 37°C with 5% CO_2_. The cell aggregates were harvested with PBS, condensed with centrifugation, and cultured in a new 60 mm non-treated culture dish. The cell culture media was changed once in 48 h up to 7 days. The microscopic images were taken in suspension culture every 48 h, and the ImageJ software collected the size of the cellular spheroids.

### 2.5 Cellular Spheroid Characterization and Drug Testing

For the field-emission scanning electron microscopy (FESEM), the spheroids were treated with 1 mL of 2% paraformaldehyde (20 mL^-1^) for 1.5 h and washed with 1 mL of PBS three times. To dehydrate the spheroids, gradual ethanol concentrations (*e*.*g*., 25, 50, 70, 90, 100, 100, and 100%) were prepared and treated to the spheroids for 20 min each. The dehydrated spheroids were deposited onto the aluminum specimen mounts (75210; Electron Microscopy Sciences, PA) and dried overnight. The samples were coated with 3.5 nm Iridium by Cressington Vacuum Coating Systems 208HRD (Ted Pella International Company, CA) and imaged under 10.0 kV with a field-emission scanning electron microscopy (Magellan 400, FEI Company, OR).

For the Live/Dead assay, the LIVE/DEAD™ Cell Imaging Kit (488/570) (Thermofisher Scientific, MA) was used following the manufacturer’s instructions. 1 mL of spheroid suspensions (*e*.*g*., 3-day-cultured and 10-day-cultured spheroids; 20 mL^-1^) were mixed with the staining solution containing both live cell indicator and dead cell indicator for 3 h followed by the 3 h fixation with 2% paraformaldehyde. The samples were kept in 2 mL PBS before imaging with A1R-MP Laser Scanning Confocal Microscopy (CLSM; Nikon, Tokyo, Japan). The images were processed for the maximum intensity projection using NIS-Elements software (Nikon, Tokyo, Japan).

For DOX dosage-dependent test, a range of DOX solutions was prepared (*e*.*g*., 0, 20, 40, 80, 160, and 320 μM) in fresh cell culture media with a final volume of 1 mL. The 10-day-cultured cellular spheroids were incubated with the DOX-embedding media for 6 h (*e*.*g*., 20 mL^-1^). Following two-time PBS washing, the spheroids were incubated with fresh cell culture media for 24 h. The spheroids were stained with a LIVE/DEAD™ Cell Imaging Kit and fixed with 2% paraformaldehyde using the same protocol and procedure as the Live/Dead assay. The microscopic images of cellular spheroids were collected using bright field microscopy for each concentration (*e*.*g*., 6-h treatment of 0, 20, 40, 80, 160, and 320 μM DOX, respectively, followed by 24-h incubation with fresh cell culture media) and for each incubation time (*e*.*g*., 6-h treatment of 120 μM DOX followed by 24, 72, 120, and 144 h of incubation with fresh cell culture media) to estimate the size change of cellular spheroids using the ImageJ software. For DOX-loaded GQD drug effect assay, *L*-GQD-DOX or *D*-GQD-DOX solutions (*e*.*g*., Prepared as GQD:DOX=0.5 mg·mL^-1^:350 μM; See **Materials and Methods Section 2.2**) were mixed with fresh cell culture media with the final concentration of DOX as 120 μM with the final volume of 1 mL. The 10-day-cultured cellular spheroids were incubated with the *L/D*-GQD-DOX-containing media for 6 h (*e*.*g*., 20 mL^-1^). Following two-time PBS washing, the spheroids were incubated with fresh cell culture media for 24 h. The spheroids were collected to perform two experiments: (1) luminescence-based cell viability assay and (2) fluorescence-based live/dead cell imaging. For luminescence-based cell viability assay, the spheroids were in a 96-well plate (*e*.*g*., 1 well^-1^ with 100 μL fresh cell culture media) and mixed with 100 μL reaction solution of CellTiter-Glo 3D cell viability assay kit (G9681; Promega, WI). The 96-well plate was gently rocked for 5 min and stabilized at room temperature for 30 min, following the manufacturer’s instruction. The luminescence from the 96-well plate was measured with Tecan Infinite 200Pro (Tecan, Männedorf, Switzerland) and normalized with the control group. For fluorescence imaging, the spheroids were stained with a LIVE/DEAD™ Cell Imaging Kit and fixed with 2% paraformaldehyde using the same protocol and procedure as the Live/Dead assay.

### 2.6 Monitoring GQD Transport into Cellular Spheroid

The effect of GQDs on early ECM was evaluated by treating *L/D*-GQDs to HepG2 aggregates (*e*.*g*., 3-day-cultured HepG2 cellular aggregates without suspension culture) or HepG2 spheroids (*e*.*g*., 10-day-cultured HepG2 cellular spheroids after suspension culture) under CLSM imaging. The 3-day-cultured cellular aggregate suspension (*e*.*g*., 20 mL^-1^; 0.9 mL) or the 10-day-cultured cellular spheroid suspension (*e*.*g*., 20 mL^-1^; 0.9 mL) and *L/D*-GQDs (1 mg·mL^-1^) were prepared. As GQDs were treated to cellular aggregates/spheroids, the images were collected under Capturing Time Series Images function (*e*.*g*., 12 recurring cycles; 1 cycle refers to Capturing *Z* Series Images for aggregates). The bright-field images, including respective temporal information, were collected throughout the aggregate region. The size distribution of aggregates/spheroids was collected using ImageJ software from bright-field channel images for each time point.

GQD transports into 3-day-cultured cellular aggregates and 10-day-cultured cellular spheroids were evaluated by CLSM imaging. The 3-day-cultured cellular aggregates, the 10-day-cultured cellular spheroid suspension (*e*.*g*., 20 mL^-1^; 0.9 mL), and *L/D*-GQDs (1 mg·mL^-1^; 0.1 mL) were prepared. Under Capturing Time Series Images function (*e*.*g*., 12 recurring cycles; 1 cycle refers to Capturing *Z* Series Images for aggregates/spheroids) on the A1R-MP CLSM, 100 μL of GQD solutions were treated gently into 15 mm glass bottom dish (801002, NEST Biotech, Wuxi, China) embedding 0.9 mL of the aggregate/spheroid suspension (*e*.*g*., Total 1 mL; 18 aggregates/spheroids; 0.1 mg GQDs). The total GQD intensity within cellular aggregates and spheroids was collected from the maximum intensity projection images from NIS-Elements software and estimated with ImageJ software. The time-lapse spatiotemporal information of the GQD transport was collected and processed with the maximum intensity projection using NIS-Elements software. ImageJ software collected and quantified the spatiotemporal information from GQD-channel images for respective time points, processing them to select the spheroid outline and quantify GQD transport.

### 2.7 Image Processing and Quantification

The oversaturation in image contrast followed by color inversion made the bright-field images into the masking layer for isolating the GQD signals only for the spheroid region. The outlined GQD channel images in the spheroid regions were quantified with further image processing for three approaches: Integrated intensity analysis, radially-averaged intensities, and time-dependent intensities at a fixed radial position. First, the integrated intensity analysis was performed with the following equation (2), where *I(x,y)* is the intensity of the GQD signal in *x* and *y* coordinates inside the spheroid region, *S* is the set of points inside the outlined region of a cellular spheroid, and *A*_*s*_ is an area of the cellular spheroid. Second, the radially-averaged intensities of the *L/D*-GQDs were quantified with the spheroid region-masked GQD channel images by using Radial Profile Angle plugin in ImageJ software. Third, the time-dependent intensities at a fixed radial position were collected from the radially-averaged intensities with fixed relative radius (*e*.*g*., 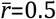 and 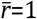 where 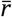 denotes relative radii to the whole spheroid radius).

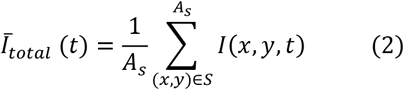

A simple diffusion equation was introduced to determine apparent diffusion coefficients, as shown in the following equation (3-4) with dimensionless parameters 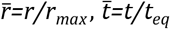 and 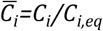, where *r*_*max*_ stands for the radius having the maximum intensity from the radially-averaged plot, *t*_*eq*_ stands for the observed time that intensity reaches a plateau, and *C*_*i,eq*_ stands for the concentration or intensity of GQDs when intensity reaches a plateau. Here, *D*_i_, the apparent diffusion coefficients of *L/D*-GQDs (*e*.*g*., *D*_L_ or *D*_D_), is a function of 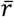 and it could be extracted by using the following equation (5), where *r*_*max*_ is spheroid radius and *t*_*eq*_ is the time that intensity reaches a plateau in the time-dependent intensities at a fixed radial position. To quantify the drug effect of DOX, the area of dead cells in the *Z*-projected CLSM image was collected and normalized with those of live area from each spheroid (*e*.*g*., extracted from the Calcein-AM channel).

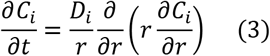

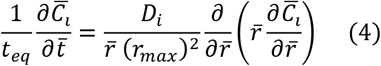

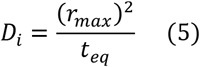

## 3. Result and Discussion

Chiral GQDs were derived by the same procedure as our previous work (*i*.*e*., EDC/NHS reaction) (Suzuki et al., 2016) from as-synthesized GQDs by a modified Hummers method (Peng et al., 2012). In brief, the edges of as-synthesized GQDs were functionalized with *L*-cysteines (*i*.*e*., *L-*GQDs) or *D*-cysteines (*i*.*e*., *D*-GQDs) under EDC/NHS reaction, aiming the amide bond between the carboxylic acid group on GQDs (*i*.*e*., -COOH) and amine group in both cysteines (*i*.*e*., -NH_2_) (Suzuki et al., 2016; Zhu et al., 2022a; Zhang et al., 2023). The sizes of *L*-GQDs and *D*-GQDs are similar to as-synthesized GQDs (**Figure 2A** and **Supplementary Figure 1A**). The frequency distribution analysis for the TEM images showed that the average size of *L*-GQDs was 7.88 ± 2.11 nm, while *D*-GQDs was 7.99 ± 2.03 nm (**Figure 2B** and **Supplementary Figure 1B**), which confirmed that *L*-GQDs and *D*-GQDs had a similar size. The FTIR analysis (**Figure 2C**) verified the chemical modification of the *L/D*-GQDs compared to the unmodified GQDs. Both *L*- and *D*-GQDs have the featured bonds in as-synthesized *L/D*-GQDs (*i*.*e*., 3400, 1409, and 1350 cm^-1^ for -OH group; 1712 cm^-1^ for -C=O group; 1602 cm^-1^ for C=C group; 1119 cm^-1^ for C-O group), as well as the *L/D*-cysteines related peak at 1250 cm^-1^ (*i*.*e*., C-N group) (**Supplementary Table 1**). This result demonstrated the successful chiral modification of GQDs with *L/D*-cysteines. CD spectra of *L/D*-GQDs displayed ellipticity peaked at the same wavelengths of 213 nm and 258 nm with opposite signs **(Figure 2D**) without significant differences in absorbance (**Supplementary Figure 2**), confirming the successful chiral modification of GQDs. Excited at 365 nm wavelength, both *L/D*-GQDs showed significant emission at ∼500 nm, regardless of the chiral modification on the particles (**Figure 2E**).

**Figure 2.**
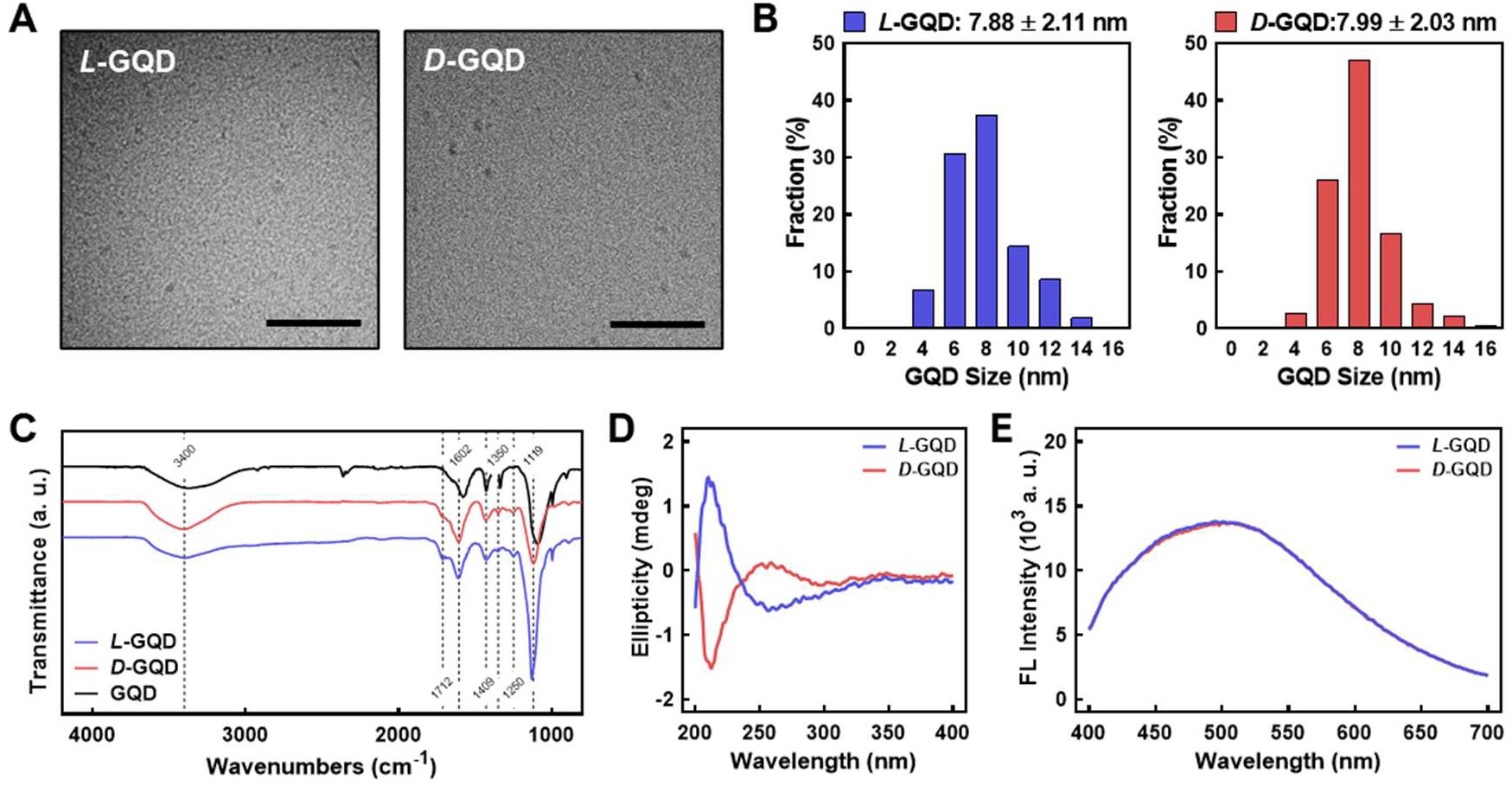
Characterizations of L/D-GQDs. **(A)** Transmission electron microscopic (TEM) images of *L*-GQDs (left) and *D*-GQDs (right) (Scale bar: 100 nm). **(B)** Size distributions of *L*-GQDs (left) and *D*-GQDs (right). The distributions were collected and analyzed from TEM images. **(C)** Fourier-transform infrared (FTIR) spectra of *L/D*-GQDs compared to the unmodified GQDs (blue: *L*-GQDs, red: *D*-GQDs, black: GQDs). **(D)** Circular Dichroism (CD) spectra of *L/D*-GQDs (blue: *L*-GQDs, red: *D*-GQDs). **(E)** Emitted fluorescent intensity of *L/D*-GQDs excited at 365 nm (blue: *L*-GQDs, red: *D*-GQDs).

To investigate the transport of chiral GQDs in cancerous tissues, tumor-like cellular spheroids were derived from HepG2 cells using the Hanging-drop method for three days (*i*.*e*., immature cellular aggregates) (Timmins and Nielsen, 2007) followed by the suspension culture for seven days (*i*.*e*., mature cellular spheroids) (Ryu et al., 2019). The serial microscopic imaging of the spheroids showed the proliferation of the cells in spheroids under the suspension culture (**Figure 3A**). The diameter of the spheroids was around 170 μm on Day 3 (*i*.*e*., cellular aggregates using the Hanging-drop method) while showing linear growth in suspension culture step up to 400 μm until Day 11 (*i*.*e*., cellular spheroids; **Figure 3B**). This result implies an optimal growth rate of the cells inside the spheroids. Under FESEM images (**Figure 3C**), characteristic comparisons between immature cellular aggregates and mature cellular spheroids were performed by observing ECM structure on the surface of cellular spheroids. Mature cellular spheroids (*i*.*e*., 10-day-cultured cellular spheroids with 7-day suspension culture after 3-day Hanging drop culture) showed denser polymeric structure on their surface than immature cellular aggregates (*i*.*e*., 3-day-cultured cellular aggregates derived from 3-day Hanging drop culture), implying the formation of more ECM that mimic the native microenvironment in matured cellular spheroids. Moreover, cellular aggregates and spheroids showed a high ratio of live cells (*e*.*g*., the green signal from Calcein-AM) to dead cells (*e*.*g*., the red signal from BOBO-3 Iodide) using Live/Dead assay, demonstrating their high cell viability (**Figure 3D**). A common chemotherapy drug, DOX, was administered into cellular spheroids to determine the dose-dependent effect of DOX on cellular spheroids. CLSM images of HepG2 spheroids stained with the Live/Dead assay (**Figure 3E**) showed an increasing portion of dead cells (*i*.*e*., Red signal from BOBO-3 iodide) in cellular spheroids as DOX concentrations increased. This result was further analyzed by the area quantification from image processing (**Figure 3F**), demonstrating a DOX dosage-responsive effect on cellular spheroids. A Fitting Hill Function estimated the effective dose at 50% death (ED_50_) as 65.29 μM, similar to the previous test of the dose-responsive effect of DOX on HepG2 cellular spheroids (Zhu et al., 2022b). Accordingly, the size of the cellular spheroids decreased as the DOX dosage increased, showing 20% shrinkage of their diameter with the 320 μM DOX treatment and 24-h incubation (**Supplementary Figure 3A**). Moreover, increasing incubation time after the DOX treatment resulted in further size decrease of cellular spheroids, implying the effective drug response (**Supplementary Figure 3B**). These results indicated the cellular spheroids are a reliable 3D culture model *in vitro* for evaluation of nanocarriers.

**Figure 3.**
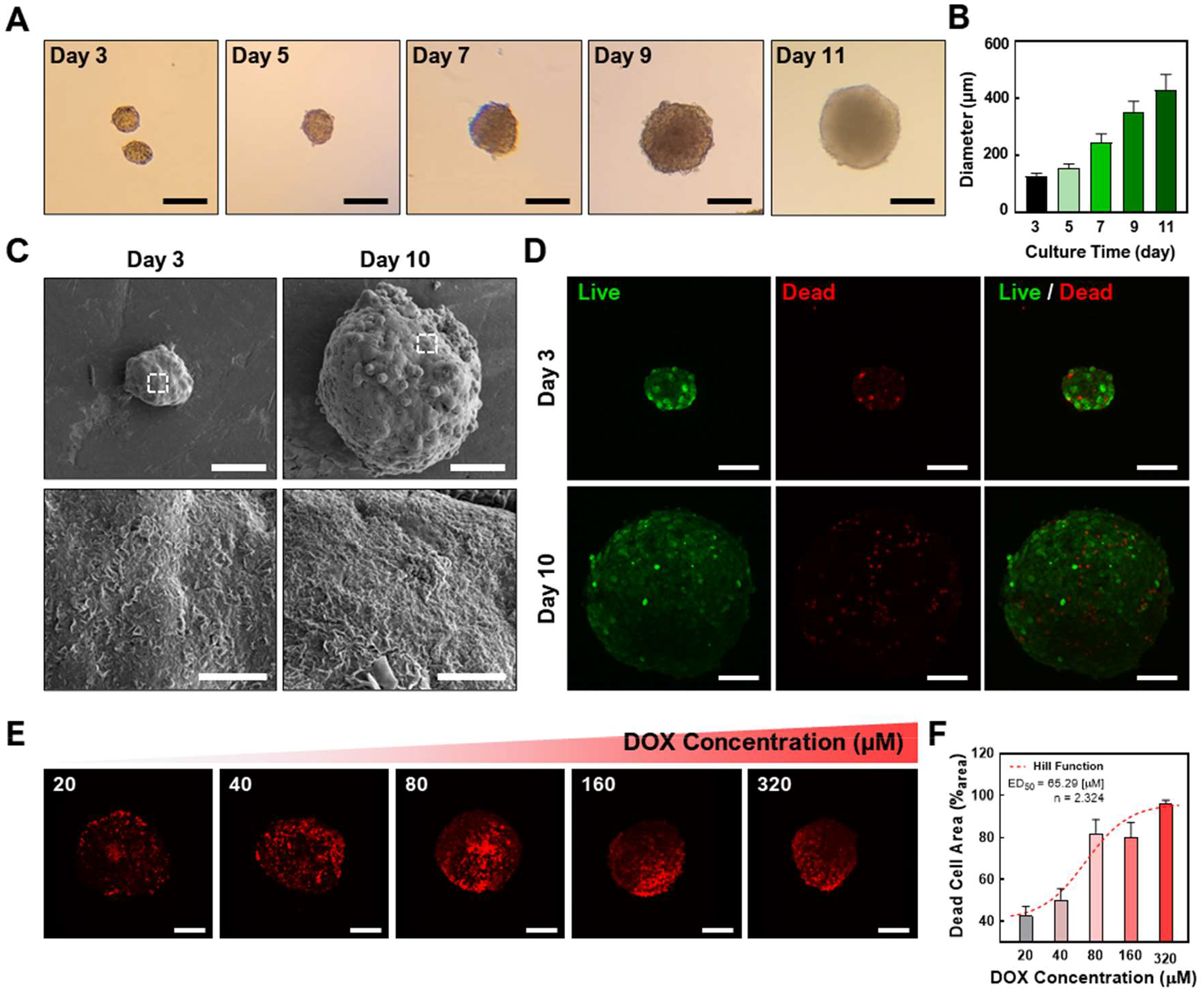
Characterizations of cellular spheroids. **(A)** Time-lapse microscopic images of the cellular spheroids in suspension culture (Scale bar: 200 μm) and **(B)** sizes of the cellular spheroid at each time point. **(C)** Field-emission scanning electron microscopic (FESEM) images of the cellular aggregates and cellular spheroids. **(D)** Confocal laser scanning microscopic (CLSM) images showing the cellular aggregates and spheroids with Live/Dead staining (Green: Calcein-AM, live cell indicator; Red channel: BOBO-3 Iodide, dead cell indicator; Scale bar: 100 μm; *Z*-projected maximum intensity images). (**E**) CLSM images showing a DOX dosage-dependent treatment in cellular spheroids. The dead cells in cellular spheroids were stained with BOBO-3 Iodide (Scale bar: 200 μm; *Z*-projected maximum intensity images). (**F**) Quantitative analysis of DOX concentration-dependent response of cellular spheroids. A Hill Function was plotted to determine ED_50_ (65.29 μM).

To investigate how the chirality of nanocarrier chiral GQDs impacts tumor tissue, we added *L/D*-GQDs into the immature/mature HepG2 cellular aggregates/spheroids (*e*.*g*., 3/10-day-cultured) and observed their size change using CLSM time-lapse imaging (**Figure 4A-B**). As a result, *L*-GQDs induced significant swelling of cellular aggregates within 20 min, while the size of cellular aggregates treated by *D***-**GQDs did not change significantly (**Figure 4A**). The size estimation by quantitative analysis revealed that *L-*GQDs-driven swelling of cellular aggregates was about a 20% increase in diameter (**Figure 4B**). In comparison, only a 2.5% diameter change was observed for *D*-GQD-treated cellular aggregates, indicating that *L*-GQDs induced a 1.41-fold structural volume change compared to *D*-GQDs. This phenomenon indicated the different chirality of GQDs induced distinct interaction between GQDs and ECM of cellular aggregates, consistent with the integrated intensity analysis (See **Supplementary Result Section 1.1**; **Supplementary Figure 4**). In contrast, mature cellular spheroids showed no significant size change after being treated with chiral GQDs (**Figure 4C, D**). These results implied chiral GQDs interacted distinctively with the immature ECM structure in cell aggregates, while not affecting the structural stability of the cellular spheroids due to the complete formation of ECM (See **Figure 3C**).

**Figure 4.**
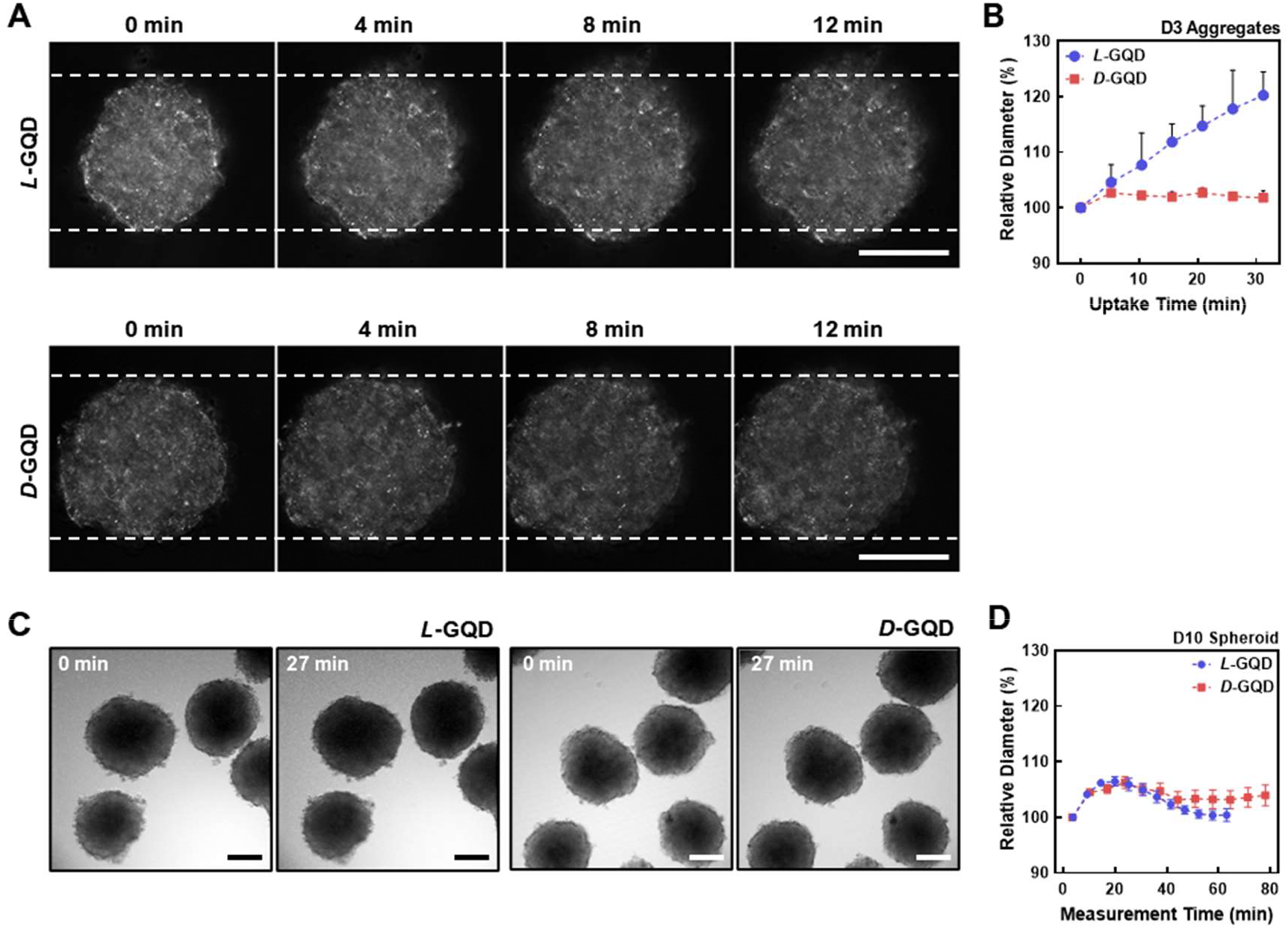
The effects of chiral GQDs on the swelling of cellular aggregates and spheroids. The bright field images were taken with CLSM under the chiral GQD treatment to cellular aggregates and spheroids. **(A)** Bright-field images showing size changes of the cellular aggregates under the *L/D*-GQD treatment (*L*-GQDs: top, *D*-GQDs: bottom, Scale bar: 100 μm). White dashed lines indicated the initial width of aggregates. **(B)** Size changes of the spheroids under the *L/D*-GQD treatments (*L*-GQDs: blue curve, *D*-GQDs: red curve; *N*=5). **(C)** Bright-field images showing the size change of the cellular spheroids under the *L/D*-GQD treatment (*L*-GQDs: left, *D*-GQDs: right; Scale bar: 100 μm). **(D)** Size changes of the cellular spheroids under the *L/D*-GQD treatment (*L*-GQDs: blue curve, *D*-GQDs: red curve; *N*=5).

To study the effect of nanocarrier’s chirality on their transport into cancerous tissue, we monitored the distribution of chiral GQDs in tumor-like cellular spheroids (*e*.*g*., 10-day-cultured cellular spheroids) throughout the time-lapse CLSM images (*i*.*e*., TRITC-filtered red channel). We continuously image the process with a 12-step protocol starting from 0 min (*i*.*e*., after adding *L/D-*GQDs to cellular spheroids), spanning less than 80 min. The GQD signals in the fluorescent images (**Figure 5A** and **Supplementary Figure 5A-B**) showed significant changes in the spheroid region as a function of time. The interface of the spheroids and solutions were distinguishable right after adding GQDs (*i*.*e*., 0 min images in **Figure 5A** and **Supplementary Figure 5A-B**), while the interface between the spheroid and the solution turned into red signals gradually over time (*i*.*e*., > 0 min images in **Figure 5A** and **Supplementary Figure 5A-B**). This phenomenon demonstrated that both *L*- and *D*-GQDs diffused into the cellular spheroid. Notably, GQD diffusion mainly occurred at the very early stage of the treatment (*e*.*g*., Intensities in the spheroid region changed between 0 min and 27 min images in **Figure 5A**) and reached a plateau later than 40 min images (See **Supplementary Figure 5A-B**).

**Figure 5.**
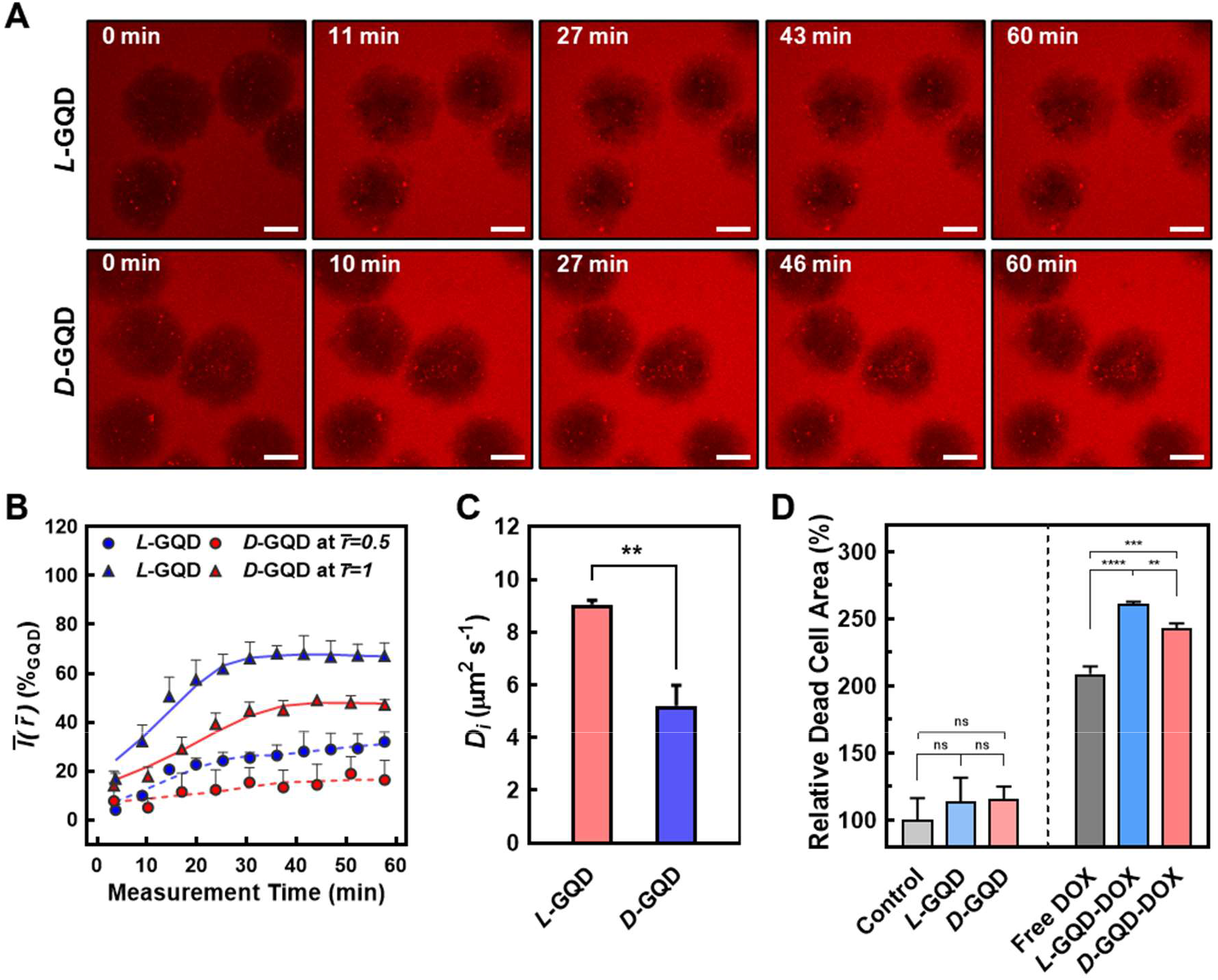
The comparison of chiral GQD diffusion to cellular spheroids. **(A)** Time-lapse CLSM images showing the transport of the GQDs into the spheroids (Red channel: GQDs; *L*-GQDs: top, *D*-GQDs: bottom; Scale bar: 100 μm; *Z*-projected maximum intensity images). **(B)** Time-dependent intensity plots at a fixed radial position. Solid lines showed the intensities in maximum radius 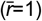 while dotted lines showed the intensities in half radius 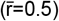. **(C)** Evaluated apparent diffusion coefficients for *L/D*-GQDs (**: *P*<0.01). **(D)** Quantitative analysis of dead cell area under DOX-load *L/D*-GQD treatment to cellular spheroids (ns: not significant, **: *P*<0.01, ***: *P*<0.001, ****: *P*<0.0001).

To quantitatively study the transport of chiral GQDs in cellular spheroids (**Figure 5B-C**), we further analyzed the GQD distribution over time by image processing (**Supplementary Figure 6**) based on the CLSM images (See **Figure 5A** and **Supplementary Figure 5A-B**; 10-day-cultured cellular spheroids). The radially averaged intensity plots were formed by averaging the intensity of GQDs from the radius of each spheroid, facilitating the GQD-signal analysis with spatial and temporal coordinates (*i*.*e*., Ī*(r,t)*; See **Materials and Methods Section 2.7**) (**Supplementary Figure 7**). Based on the analysis, *L*-GQDs exhibited 1.5-fold higher plateau intensities than *D*-GQDs at the edge of the cellular spheroid while showing a shorter time to reach a plateau at *r*_*max*_ than *D*-GQDs. This result implies that *L-*GQDs diffuse into the cellular spheroids more effectively and efficiently than *D*-GQDs, consistent with the integrated intensity analysis (See **Supplementary Result Section 1.1**; **Supplementary Figure 8**).

To evaluate the chirality-induced effect of GQD transport precisely, time-dependent intensities at a fixed radial position were collected from the spatiotemporal distribution (See **Supplementary Figure 7**) and represented with relative radii: the outer shell of the spheroids (*i*.*e*.,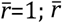 denotes relative radii to the whole spheroid radius) and half of the radius (*i*.*e*.,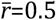) (**Figure 5B**). The plots from the outer shell showed the intensity saturation at specific times, *t*_*eq*_ (*i*.*e*.,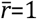 for both *L*-GQDs and *D*-GQDs; solid lines in **Figure 5B**), while intensities from the half radius were increasing throughout the measurement (*i*.*e*.,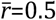 for both *L-*GQDs and *D*-GQDs; dotted lines in **Figure 5B**). The *t*_*eq*_ values were estimated as 22.6 ± 1.2 min for *L*-GQDs and 31.4 ± 3.7 min for *D*-GQDs, implying *L*-GQDs have a higher apparent diffusion coefficient (*i*.*e*., *D*_*i*_) than *D*-GQDs. These estimations suggested that the transport of *L*-GQDs in cellular spheroids is much faster than that of *D*-GQDs.

To estimate the *D*_*i*_ values of *L/D*-GQDs in tumor-like tissue (*i*.*e*., *D*_*L*_ or *D*_*D*_), a simple diffusion model was introduced as shown in the equation (3-4) (See **Materials and Methods Section 2.7**). The 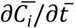 term is negligible (*i*.*e*.,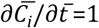 Due to the dimensionless form) and it can be assumed that the radially-averaged plots are linear (*i*.*e*., 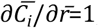; See **Supplementary Figure 7**). Hence, *D*_*i*_ is the function of 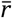 and could be extracted by using equation (5): *D*_*i*_*=*(*r*_*max*_)^*2*^*/t*_*eq*_, where *i* denotes *L* or *D* for *L/D*-GQDs (*i*.*e*., *D*_*L*_ or *D*_*D*_). Extracted *D*_*L*_ was 9.04 μm^2^·s^-1^ and *D*_*D*_ was 5.20 μm^2^·s^-1^ (**Figure 5C**), showing statistically significant differences. Notably, *D*_*L*_ was 1.7-fold higher than *D*_*D*_, indicating again that *L*-GQDs are more favorable to diffuse into the cellular spheroid. Interestingly, the values of *D*_*L*_ and *D*_*D*_ are much smaller than the diffusivity calculated from the Stokes-Einstein equation (*e*.*g*., 62.3 μm^2^·s^-1^ for 7 nm NP dispersed in water at 25°C) (Rudyak, 2016), similar to the diffusivity of 44 ± 1 nm-sized NPs (*e*.*g*., 7.31 ± 0.4 μm^2^·s^-1^ dispersed in water at 25°C) (Mun et al., 2014). This fact implies NP diffusion was hindered by the densely packed ECM structures of cancerous cellular spheroids. In summary, our studies on chiral GQD diffusion through cellular spheroids showed enhanced transport of *L*-GQDs compared to those of *D*-GQDs, implying chiral modification of NPs can enhance their diffusion and potentially improve drug delivery efficiency in tumor-like tissue.

As a proof-of-concept, we employed chiral GQDs as nanocarriers for small chemotherapy drugs (*e*.*g*., DOX) and investigated their delivery efficiency into tumor-like tissue. Due to its planar anthraquinone structure, chiral GQDs can be loaded with DOX *via* the π-π stacking (**Supplementary Figure 9**) (Liu et al., 2009; Wo et al., 2016; Askari et al., 2021; Sawy et al., 2021; Kiani Nejad et al., 2022; Mirzaei-Kalar et al., 2022). Two drug-loaded nanocarriers, *L*-GQD-DOX and *D*-GQD-DOX, were tested in mature cellular spheroids and compared to the controls (*e*.*g*., Non-treated, *L*-GQD-treated, and *D*-GQD-treated groups as negative controls; free DOX-treated group as a positive control). As a result, both nanocarrier groups loaded with DOX (*i*.*e*., *L*-GQD-DOX and *D*-GQD-DOX groups) show significant increases in the ratio of dead cells than all negative controls in the CLSM images with Live/Dead staining (**Supplementary Figure 10** and **Figure 5D**). Compared to the positive control, free DOX group, the *L/D*-GQD-DOX groups showed a higher ratio of dead cells: 25% increase with *L*-GQD-DOX and 16% increase with *D*-GQD-DOX showing a statistical significance (**Figure 5D**). Moreover, *L*-GQD-induced enhanced delivery of DOX led to a lower living cell activity as the cell viability in **Supplementary Figure 11**, showing a statistical significance between free DOX and *L*-GQD-DOX groups. Combining all, these facts demonstrates that the chirality of nanocarriers can improve the delivery efficiency of drugs by enhanced transport through ECM.

## 4. Conclusion

Our study successfully developed chiral nanocarriers, *L*- and *D*-GQDs and evaluated their transport in tumor-like cellular spheroids. We discovered that left-handed chiral nanocarrier, *L*-GQDs, induced structural swelling of cellular aggregates with 15% in their diameter while right-handed nanocarrier (*D*-GQDs) showed limited effect, indicating the distinct interactions of the chiral GQDs with the immature ECM structure. Moreover, we demonstrated the impact of nanocarrier chirality on its transport and diffusion in tumor-like cellular spheroids: a 1.7-fold higher *D*_*i*_ of *L*-GQDs than that of *D*-GQDs. Furthermore, we conducted a proof-of-concept design for enhanced DOX delivery, *L*-GQD-DOX, based on the unique feature of *L*-GQDs, leading to a 25% increase in drug effect compared to free DOX. Overall, our work indicated the importance to consider chirality when designing drug carriers, as it is promising to enhance the transport and delivery of small drugs in 3D tumor-like tissue.

## 6. Author Contributions

HJ and YW conceptualized and designed the study and all the experiments. YW supervised overall conceptualization, validation, and data analysis. RZ synthesized and characterized chiral GQDs. GK synthesized and characterized DOX-loaded chiral GQDs. HJ performed cell culture-related works and fabricated 3D cellular spheroids. HJ performed cell-related imaging and image processing. HJ wrote the original draft, and YW supervised all the work. All authors discussed the results and commented on the manuscript.

## 8. Acknowledgements

All the CLSM imaging was carried out in part in the Notre Dame Integrated Imaging Facility, University of Notre Dame, using A1R-MP Laser Scanning Confocal Microscopy. We thank Sara Cole for the knowledge and expertise as well as time towards this research.

## References

Al-Hajaj, N. A., Moquin, A., Neibert, K. D., Soliman, G. M., Win nik, F. M., and Maysinger, D. (2011). Short Ligands Affect Modes of QD Uptake and Elimination in Human Cell s. ACS Nano 5, 4909–4918. doi: 10.1021/nn201009w.

Askari, E., Naghib, S. M., Zahedi, A., Seyfoori, A., Zare, Y., and Rhee, K. Y. (2021). Local Delivery of Chemotherapeutic Agent in Tissue Engineering Based on Gelatin/graph ene Hydrogel. J Mater Res 12, 412–422. doi: 10.1016/j.jmrt.2021.02.084.

Biswas, M. C., Islam, M. T., Nandy, P. K., and Hossain, M. M. (2021). Graphene Quantum Dots (GQDs) for Bioimaging and Drug Delivery Applications: A Review. ACS Mater Lett. doi: 10.1021/acsmaterialslett.0c00550.

Charoen, K. M., Fallica, B., Colson, Y. L., Zaman, M. H., and Grinstaff, M. W. (2014). Embedded Multicellular Spheroids as a Biomimetic 3D Cancer Model for Evaluating Drug and Drug-Device Combinations. Biomater. 35, 2264 –2271. doi: 10.1016/j.biomaterials.2013.11.038.

Chauhan, V. P., Lanning, R. M., Diop-Frimpong, B., Mok, W., Brown, E. B., Padera, T. P., et al. (2009). Multiscale Mea surements Distinguish Cellular and Interstitial Hindrances to Diffusion In Vivo. Biophys J 97, 330–336. doi: 10.1016/j.bpj.2009.03.064.

Chen, C. S. (2016). 3D Biomimetic Cultures: The Next Platform for Cell Biology. Trends Cell Biol 26, 798–800. doi: 10.1016/j.tcb.2016.08.008.

Cooper, G. M. (2000). “Transport of Small Molecules,” in Th e Cell: A Molecular Approach. 2nd edition (Sinauer Associates). Available at: https://www.ncbi.nlm.nih.gov/books/NBK9847/ [Accessed April 6, 2023].

Döring, A., Ushakova, E., and Rogach, A. L. (2022). Chiral Carbon Dots: Synthesis, Optical Properties, and Emerging Applications. Light Sci Appl 11, 75. doi: 10.1038/s41377-022-00764-1.

Du, X., Zhou, J., Wang, J., Zhou, R., and Xu, B. (2017). Chirality Controls Reaction-Diffusion of Nanoparticles for Inhi biting Cancer Cells. ChemNanoMat 3, 17–21. doi: 10.1002/cnma.201600258.

Du, Z., Liu, C., Song, H., Scott, P., Liu, Z., Ren, J., et al. (2020). Neutrophil-Membrane-Directed Bioorthogonal Synthe sis of Inflammation-Targeting Chiral Drugs. Chem 6, 20 60–2072. doi: 10.1016/j.chempr.2020.06.002.

Fulaz, S., Vitale, S., Quinn, L., and Casey, E. (2019). Nanoparticle–Biofilm Interactions: The Role of the EPS Matrix. Trends Microbiol 27, 915–926. doi: 10.1016/j.tim.2019.07.004.

Gao, Y., Li, M., Chen, B., Shen, Z., Guo, P., Wientjes, M. G., et a l. (2013). Predictive Models of Diffusive Nanoparticle Transport in 3-Dimensional Tumor Cell Spheroids. AA PS J 15, 816–831. doi: 10.1208/s12248-013-9478-2.

Gunti, S., Hoke, A. T. K., Vu, K. P., and London, N. R. (2021). Organoid and Spheroid Tumor Models: Techniques an d Applications. Cancers 13, 874. doi: 10.3390/cancers13040874.

Hai, X., Feng, J., Chen, X., and Wang, J. (2018). Tuning the Optical Properties of Graphene Quantum Dots for Biosen sing and Bioimaging. J Mater Chem B 6, 3219–3234. doi: 10.1039/C8TB00428E.

Henna, T. K., and Pramod, K. (2020). “Biocompatibility of Graphene Quantum Dots and Related Materials,” in Handbook of Biomaterials Biocompatibility Woodhead Publishing Series in Biomaterials., ed. M. Mozafari (Woodhead Publishing), 353–367. doi: 10.1016/B978-0-08-102967-1.00017-7.

Hsu, Y.-H., Moya, M. L., Abiri, P., Hughes, C. C. W., George, S. C., and Lee, A. P. (2012). Full Range Physiological Mass Transport Control in 3D Tissue Cultures. Lab Chip 13, 81–89. doi: 10.1039/C2LC40787F.

Huang, Y., Fu, Y., Li, M., Jiang, D., Kutyreff, C. J., Engle, J. W., et al. (2020a). Chirality-Driven Transportation and Oxid ation Prevention by Chiral Selenium Nanoparticles. Angew. Chem. 132, 4436–4444. doi: 10.1002/ange.201910615.

Huang, Y., Fu, Y., Li, M., Jiang, D., Kutyreff, C. J., Engle, J. W., e t al. (2020b). Chirality-Driven Transportation and Oxi dation Prevention by Chiral Selenium Nanoparticles. Angew Chem Int Ed Engl 59, 4406–4414. doi: 10.1002/anie.201910615.

Jiang, S., Chekini, M., Qu, Z.-B., Wang, Y., Yeltik, A., Liu, Y., et al. (2017). Chiral Ceramic Nanoparticles and Peptide C atalysis. J Am Chem Soc 139, 13701–13712. doi: 10.1021/jacs.7b01445.

Kiani Nejad, Z., Akbar Khandar, A., and Khatamian, M. (2022). Graphene Quantum Dots Based MnFe2O4@SiO2 M agnetic Nanostructure as a pH-Sensitive Fluorescence Resonance Energy Transfer (FRET) System to Enhance the Anticancer Effect of the Drug. Int J Pharm 628, 12 2254. doi: 10.1016/j.ijpharm.2022.122254.

Koomullil, R., Tehrani, B., Goliwas, K., Wang, Y., Ponnazhagan, S., Berry, J., et al. (2021). Computational Simulation of Exosome Transport in Tumor Microenvironment. Front Med 8. Available at: https://www.frontiersin.org/articles/10.3389/fmed.2021.643793 [Accessed April 6, 2023].

Leedale, J. A., Kyffin, J. A., Harding, A. L., Colley, H. E., Murdoch, C., Sharma, P., et al. (2020). Multiscale Modelling of Drug Transport and Metabolism in Liver Spheroids. Interface Focus 10, 20190041. doi: 10.1098/rsfs.2019.0041.

Lenzini, S., Bargi, R., Chung, G., and Shin, J.-W. (2020). Matrix Mechanics and Water Permeation Regulate Extracell ular Vesicle Transport. Nat Nanotechnol 15, 217–223. doi: 10.1038/s41565-020-0636-2.

Li, G., Liu, Z., Gao, W., and Tang, B. (2023). Recent Rdvance ment in Graphene Quantum Dots Based Fluorescent Sensor: Design, Construction and Bio-Medical Applications. Coord Chem Rev 478, 214966. doi: 10.1016/j.ccr.2022.214966.

Liu, Z., Fan, A. C., Rakhra, K., Sherlock, S., Goodwin, A., Chen, X., et al. (2009). Supramolecular Stacking of Doxorubicin on Carbon Nanotubes for In Vivo Cancer Therapy. Angew Chem Int Ed Engl 48, 7668–7672. doi: 10.1002/anie.200902612.

Ma, W., Xu, L., de Moura, A. F., Wu, X., Kuang, H., Xu, C., et al. (2017). Chiral Inorganic Nanostructures. Chem Rev 11 7, 8041–8093. doi: 10.1021/acs.chemrev.6b00755.

Mirzaei-Kalar, Z., Kiani Nejad, Z., and Khandar, A. A. (2022). New ZnFe2O4@SiO2@graphene Quantum Dots as an Effective Nanocarrier for Targeted DOX Delivery and C T-DNA Binder. J Mol Liq 363, 119904. doi: 10.1016/j.molliq.2022.119904.

Mun, E. A., Hannell, C., Rogers, S. E., Hole, P., Williams, A. C., and Khutoryanskiy, V. V. (2014). On the Role of Specific Interactions in the Diffusion of Nanoparticles in Aqueous Polymer Solutions. Langmuir 30, 308–317. doi: 10.1021/la4029035.

Ng, C. P., and Pun, S. H. (2008). A Perfusable 3D Cell–Matrix Tissue Culture Chamber for In Situ Evaluation of Nano particle Vehicle Penetration and Transport. Biotechnol Bioeng 99, 1490–1501. doi: 10.1002/bit.21698.

Park, H., and Park, S. Y. (2022). Enhancing the Alkaline Hydrogen Evolution Reaction of Graphene Quantum Dots by Ethylenediamine Functionalization. ACS Appl Mater Interfaces 14, 26733–26741. doi: 10.1021/acsami.2c04703.

Peng, J., Gao, W., Gupta, B. K., Liu, Z., Romero-Aburto, R., Ge, L., et al. (2012). Graphene Quantum Dots Derived from Carbon Fibers. Nano Let 12, 844–849. doi: 10.1021/nl2038979.

Peng, Z., Yuan, L., XuHong, J., Tian, H., Zhang, Y., Deng, J., et al. (2021). Chiral Nanomaterials for Tumor Therapy: Autophagy, Apoptosis, and Photothermal Ablation. J Nanobiotechnology 19, 220. doi: 10.1186/s12951-021-00965-7.

Powell, M. E., Evans, C. D., Bull, S. D., James, T. D., and Fordr ed, P. S. (2012). “Spectroscopic Analysis: Diastereomeric Derivatization for Spectroscopy,” in Comprehensive Chirality, eds. E. M. Carreira and H. Yamamoto (London, U. K.: Elsevier), 571–599. doi: 10.1016/B978-0-08-095167-6.00845-4.

Rudyak, V. Ya. (2016). “Diffusion of Nanoparticles in Gases and Liquids,” in Handbook of Nanoparticles, ed. M. Alio fkhazraei (Cham: Springer International Publishing), 1 193–1218. doi: 10.1007/978-3-319-15338-4_54.

Ryu, N.-E., Lee, S.-H., and Park, H. (2019). Spheroid Culture System Methods and Applications for Mesenchymal St em Cells. Cells 8, 1620. doi: 10.3390/cells8121620.

Salam, A. (1991). The Role of Chirality in the Origin of Life. J Mol Evol 33, 105–113. doi: 10.1007/BF02193624.

Sarasij, R. C., Mayor, S., and Rao, M. (2007). Chirality-Induce d Budding: A Raft-Mediated Mechanism for Endocytosis and Morphology of Caveolae? Biophys J 92, 3140–3158. doi: 10.1529/biophysj.106.085662.

Sattari, S., Adeli, M., Beyranvand, S., and Nemati, M. (2021). Functionalized Graphene Platforms for Anticancer Dru g Delivery. Int J Nanomedicine 16, 5955–5980. doi: 10.2147/IJN.S249712.

Sawy, A. M., Barhoum, A., Abdel Gaber, S. A., El-Hallouty, S. M., Shousha, W. G., Maarouf, A. A., et al. (2021). Insight s of Doxorubicin Loaded Graphene Quantum Dots: Syn thesis, DFT Drug Interactions, and Cytotoxicity. Mater Sci Eng C 122, 111921. doi: 10.1016/j.msec.2021.111921.

Shanker, G., Allen, J. W., Mutkus, L. A., and Aschner, M. (2001). The Uptake of Cysteine in Cultured Primary Astrocytes and Neurons. Brain Res 902, 156–163. doi: 10.1016/S0006-8993(01)02342-3.

Shanker, G., and Aschner, M. (2001). Identification and Characterization of Uptake Systems for Cystine and Cysteine in Cultured Astrocytes and Neurons: Evidence for Methylmercury-Targeted Disruption of Astrocyte Transport. J Neurosci Res 66, 998–1002. doi: 10.1002/jnr.10066.

Shao, Y., Yang, G., Lin, J., Fan, X., Guo, Y., Zhu, W., et al. (2021). Shining Light on Chiral Inorganic Nanomaterials fo r Biological Issues. Theranostics 11, 9262–9295. doi: 10.7150/thno.64511.

Sherman, I. A., and Fisher, M. M. (1986). Hepatic Transport of Fluorescent Molecules: In Vivo Studies Using Intravital TV Microscopy. Hepatology 6, 444–449. doi: 10.1002/hep.1840060321.

Speirs, A. L. (1962). Thalidomide and congenital Abnormalities. The Lancet 279, 303–305. doi: 10.1016/S0140-6736(62)91248-5.

Suzuki, N., Wang, Y., Elvati, P., Qu, Z.-B., Kim, K., Jiang, S., et al. (2016). Chiral Graphene Quantum Dots. ACS Nano 1 0, 1744–1755. doi: 10.1021/acsnano.5b06369.

Tian, P., Tang, L., Teng, K. S., and Lau, S. P. (2018). Graphene Quantum Dots from Chemistry to Applications. Mater Today Chem 10, 221–258. doi: 10.1016/j.mtchem.2018.09.007.

Timmins, N. E., and Nielsen, L. K. (2007). “Generation of Multicellular Tumor Spheroids by the Hanging-Drop Method,” in Tissue Engineering Methods in Molecular Medic ineTM., eds. H. Hauser and M. Fussenegger (Totowa, NJ: Humana Press), 141–151. doi: 10.1007/978-1-59745-443-8_8.

Vázquez-Nakagawa, M., Rodríguez-Pérez, L., Martín, N., and Herranz, M. Á. (2022). Supramolecular Assembly of E dge Functionalized Top-Down Chiral Graphene Quantum Dots. Angew Chem Int Ed Engl 61, e202211365. doi: 10.1002/anie.202211365.

Wang, S., Cole, I. S., Zhao, D., and Li, Q. (2016). The Dual Roles of Functional Groups in the Photoluminescence of Graphene Quantum Dots. Nanoscale 8, 7449–7458. doi: 10.1039/C5NR07042B.

Wang, X., Wu, B., Zhang, Y., Dou, X., Zhao, C., and Feng, C. (2022a). Polydopamine-Doped Supramolecular Chiral Hydrogels for Postoperative Tumor Recurrence Inhibition and Simultaneously Enhanced Wound Repair. Acta Biomater 153, 204–215. doi: 10.1016/j.actbio.2022.09.012.

Wang, X., Wu, B., Zhang, Y., and Feng, C. (2022b). Chiral Gra phene-Based Supramolecular Hydrogels Toward Tumor Therapy. Polym Chem 13, 1685–1694. doi: 10.1039/D1PY01724A.

Wang, Y., Bahng, J. H., Che, Q., Han, J., and Kotov, N. A. (2015). Anomalously Fast Diffusion of Targeted Carbon Nanotubes in Cellular Spheroids. ACS Nano 9, 8231–8238. doi: 10.1021/acsnano.5b02595.

Wang, Y., and Jeon, H. (2022). 3D Cell Cultures Toward Quantitative High-Throughput Drug Screening. Trends Pharmacol. Sci. 43, 569–581. doi: 10.1016/j.tips.2022.03.014.

Wang, Y., Jiang, Z., Xu, W., Yang, Y., Zhuang, X., Ding, J., et al. (2019). Chiral Polypeptide Thermogels Induce Control led Inflammatory Response as Potential Immunoadjuv ants. ACS Appl Mater Interfaces 11, 8725–8730. doi: 10.1021/acsami.9b01872.

Warning, L. A., Miandashti, A. R., McCarthy, L. A., Zhang, Q., Landes, C. F., and Link, S. (2021). Nanophotonic Appro aches for Chirality Sensing. ACS Nano 15, 15538–15566. doi: 10.1021/acsnano.1c04992.

Wo, F., Xu, R., Shao, Y., Zhang, Z., Chu, M., Shi, D., et al. (2016). A Multimodal System with Synergistic Effects of Magneto-Mechanical, Photothermal, Photodynamic and C hemo Therapies of Cancer in Graphene-Quantum Dot-Coated Hollow Magnetic Nanospheres. Theranostics 6, 485–500. doi: 10.7150/thno.13411.

Wu, W., and Pauly, M. (2022). Chiral Plasmonic Nanostructures: Recent Advances in Their Synthesis and Applications. Mater Adv 3, 186–215. doi: 10.1039/D1MA00915J.

Yao, C., Tu, Y., Ding, L., Li, C., Wang, J., Fang, H., et al. (2017). Tumor Cell-Specific Nuclear Targeting of Functionalized Graphene Quantum Dots In Vivo. Bioconjug Chem 2 8, 2608–2619. doi: 10.1021/acs.bioconjchem.7b00466.

Yeom, J., Guimaraes, P. P. G., Ahn, H. M., Jung, B.-K., Hu, Q. M cHugh, K., et al. (2020). Chiral Supraparticles for Controllable Nanomedicine. Adv Mater 32, 1903878. doi: 10.1002/adma.201903878.

Zhang, Y., Zhu, Y., Kim, G., Wang, C., Zhu, R., Lu, X., et al. (2023). Chiral Graphene Quantum Dots Enhanced Drug Loa ding into Exosomes. Acs Nano, in press. doi: 10.1021/acsnano.3c00305.

Zhao, B., Yang, S., Deng, J., and Pan, K. (2021). Chiral Graphene Hybrid Materials: Structures, Properties, and Chiral Applications. Adv Sci 8, 2003681. doi: 10.1002/advs.202003681.

Zhao, X., Zang, S.-Q., and Chen, X. (2020). Stereospecific Interactions Between Chiral Inorganic Nanomaterials and Biological Systems. Chem Soc Rev 49, 2481–2503. doi: 10.1039/D0CS00093K.

Zhu, R., Makwana, K. M., Zhang, Y., Rajewski, B. H., Valle, J. R. D., and Wang, Y. (2022a). Inhibition and Disassembly of Tau Aggregates by Engineered Graphene Quantum Dots. 2022.12.29.522245. doi: 10.1101/2022.12.29.522245.

Zhu, X., Wu, Q., He, Y., Gao, M., Li, Y., Peng, W., et al. (2022b). Fabrication of Size-Controllable and Arrangement-Or derly HepG2 Spheroids for Drug Screening via Decellularized Liver Matrix-Derived Micropattern Array Chips. ACS Omega 7, 2364–2376. doi: 10.1021/acsomega.1c06302.

Ziemys, A., Kojic, M., Milosevic, M., Schrefler, B., and Ferrari, M. (2018). Multiscale Models for Transport and Biodistribution of Therapeutics in Cancer. Comput Aided Che m Eng, 209–237. doi: 10.1016/B978-0-444-63964-6.00007-6.

