## Supplementary Material for "Chirality-Enhanced Transport and Drug Delivery of Graphene Nanocarriers to Tumor-like Cellular Spheroid"

### 1 Supplementary Result

#### 1.1 GQD Transport Estimation with Integrated Intensity Plot

To evaluate the chiral GQD transport into cellular aggregates and spheroids, we analyzed the integrated intensities of GQDs in the aggregate/spheroid regions (See **Materials and Methods Section 2.7**). For integrated intensity analysis, the GQD signal in the spheroid regions was collected and integrated throughout the spheroid area, showing the integrated intensity as a function of time (*i.e.*, $\bar{I}_{total}(t)$; See equation (1) in **Materials and Methods Section 2.7**). **Supplementary Figure 4** shows the integrated intensity of each GQD within the cellular aggregate region (*e.g.*, 3-day-cultured cellular aggregates; *N*=3). As a result, *L*-GQD showed a faster intensity increase than *D*-GQD, indicating the stronger transport of *L*-GQD within the cellular aggregates. Similarly, **Supplementary Figure 8** shows the integrated intensity of each GQD within the cellular spheroid region (*e.g.*, 10-day-cultured cellular spheroids; *N*=3). As a result, integrated intensities of *L/D*-GQDs increased over time and reached each plateau, but their values differed significantly. Additionally, the times to reach a plateau of integrated intensities of *L/D*-GQDs were distinct: *L*-GQDs reached a plateau at nearly 20 min while that of *D*-GQD was 40 min, showing different observed times to reach the plateau. These facts imply the distinct differences between *L/D*-GQDs in their transport to tumor-like tissue.

#### 2 Supplementary Figures and Tables

##### 2.1 Supplementary Figures

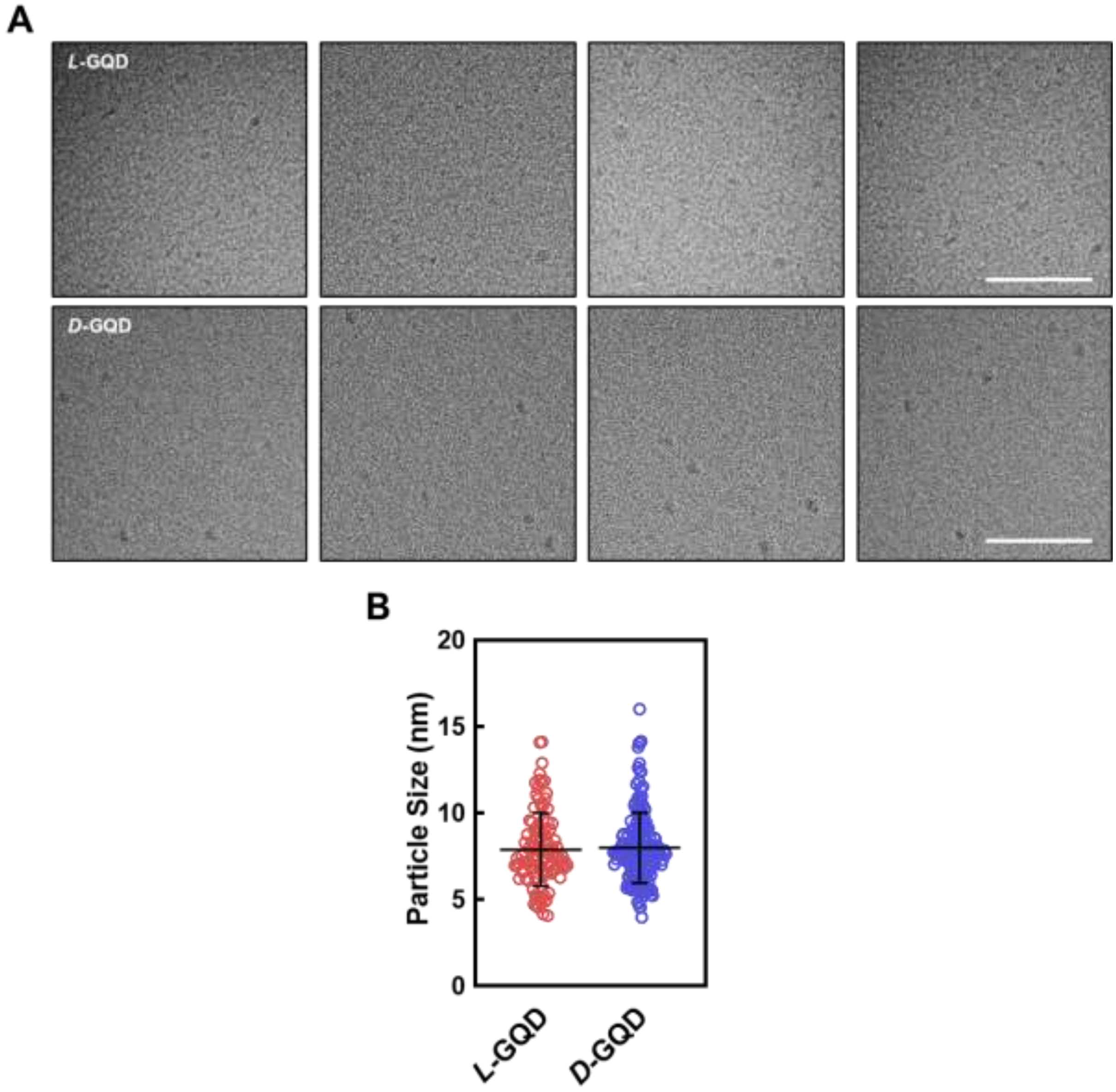


**Supplementary Figure 1. (A)** Transmission electron microscopic (TEM) images of left/right-handed graphene quantum dots (*L*/*D*-GQDs) (Top: *L*-GQDs; bottom: *D*-GQDs; Scale bar: 100 nm). **(B)** Size distribution plots for *L*/*D*-GQDs.

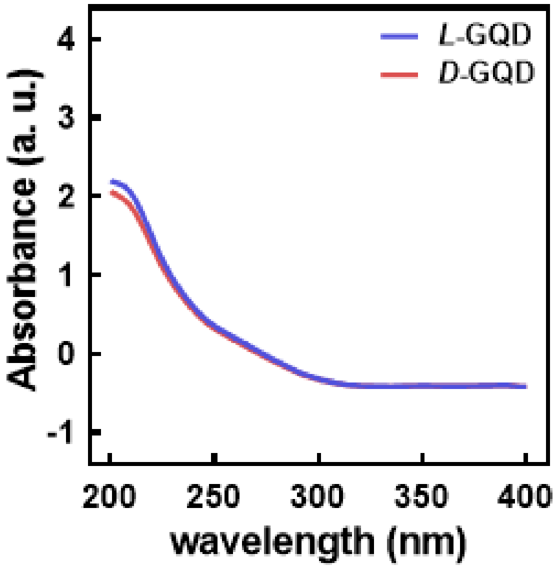


**Supplementary Figure 2.** Absorbance spectra of *L*/*D*-GQDs.

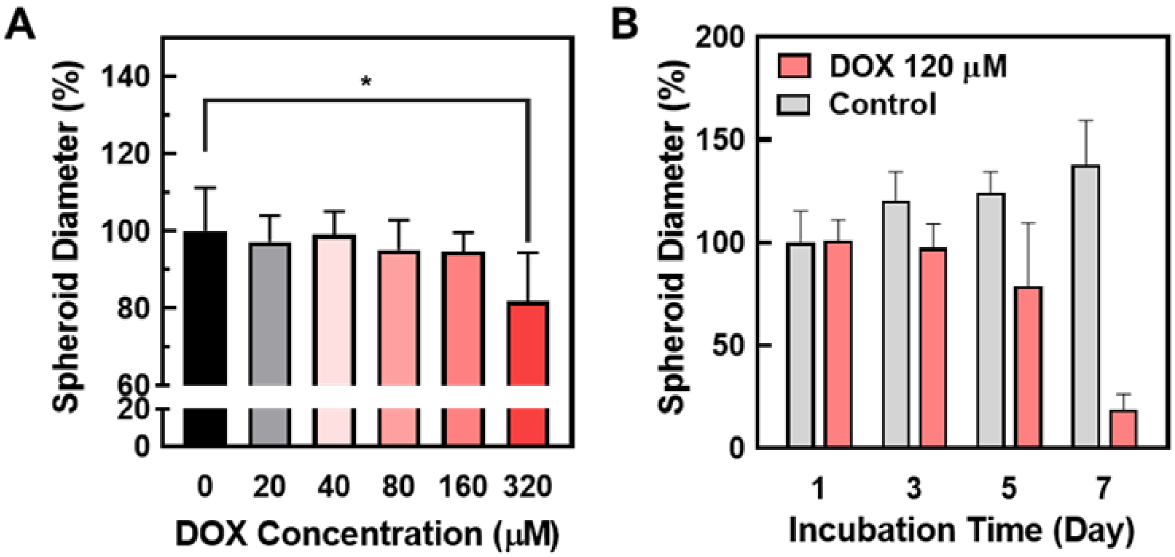


**Supplementary Figure 3.** Size changes of cellular spheroids (*e.g.*, 10-day-cultured cellular spheroids) upon doxorubicin (DOX) treatment. **(A)** DOX concentration-dependent spheroid size changes. A range of DOX was added to cellular spheroids for a 6-h treatment followed by a 24-h incubation with fresh cell culture medium (*e.g.*, 0, 20, 40, 80, 160, and 320 μM, respectively; $N$=5; *: $P$<0.05). **(B)** DOX-induced size changes of cellular spheroids upon increasing incubation time. 120 μM of DOX was added to cellular spheroids for a 6-h treatment, followed by the size measurement at respective incubation times (*e.g.*, 24, 72, 120, and 144 h; $N$=5).

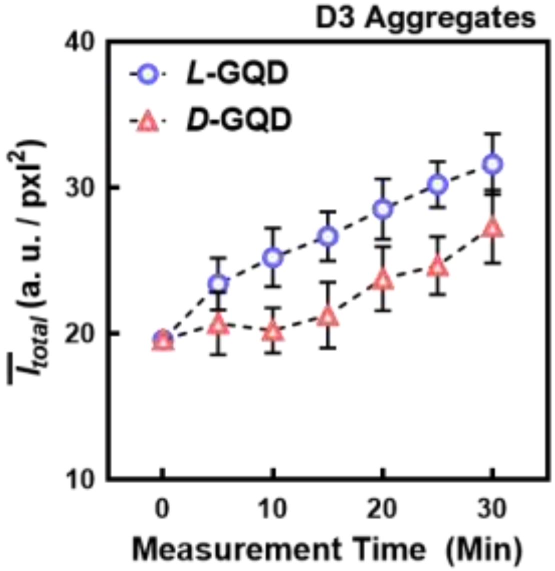


**Supplementary Figure 4.** Integrated GQD intensities increased within the region of cellular aggregates (*e.g.*, 3-day-cultured cellular aggregates) as a function of measurement time (min). The blue curve showed the result from *L*-GQD-treated aggregates, while the red curve showed the result from *D*-GQD-treated aggregates.

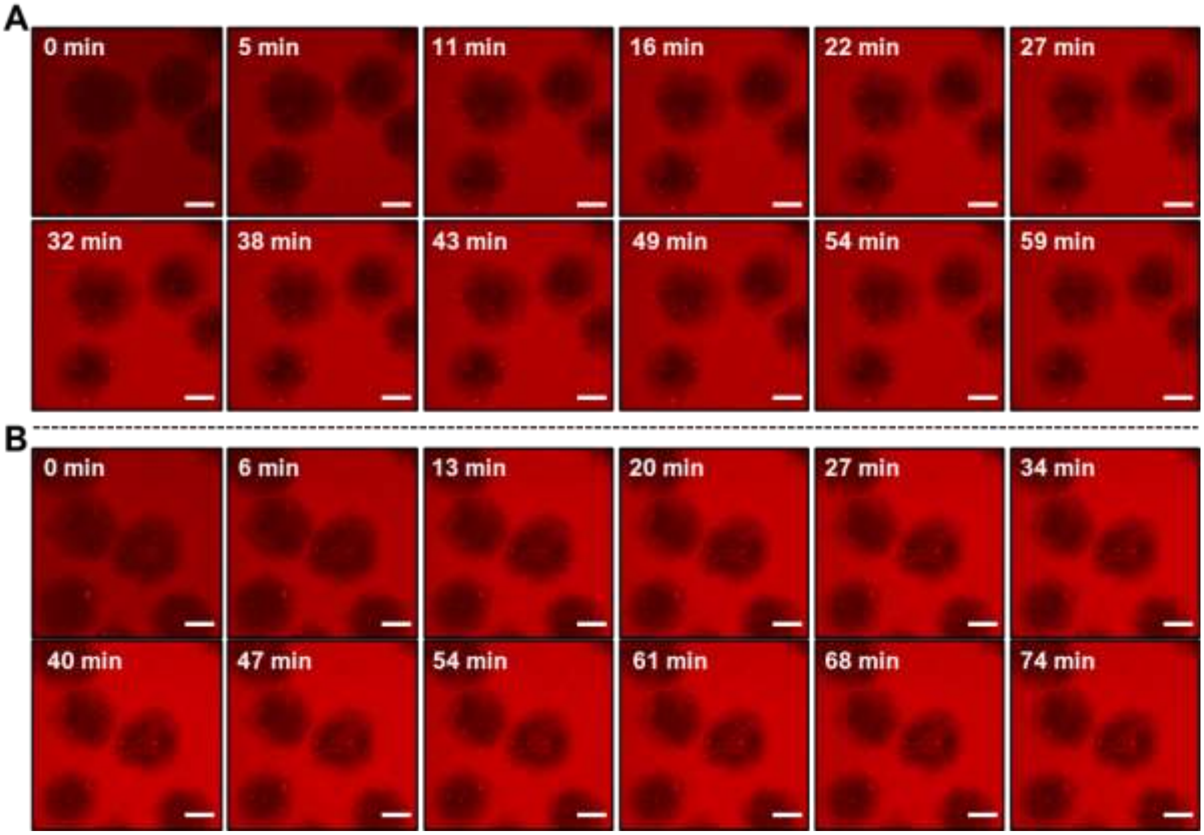


**Supplementary Figure 5.** Overall time-lapse confocal laser scanning electron microscopic (CLSM) images of GQD channel for *L/D*-GQD treatment to cellular spheroids. **(A)** The *L*-GQD transport monitoring. **(B)** The *D*-GQD transport monitoring (Scale bar: 200 µm).

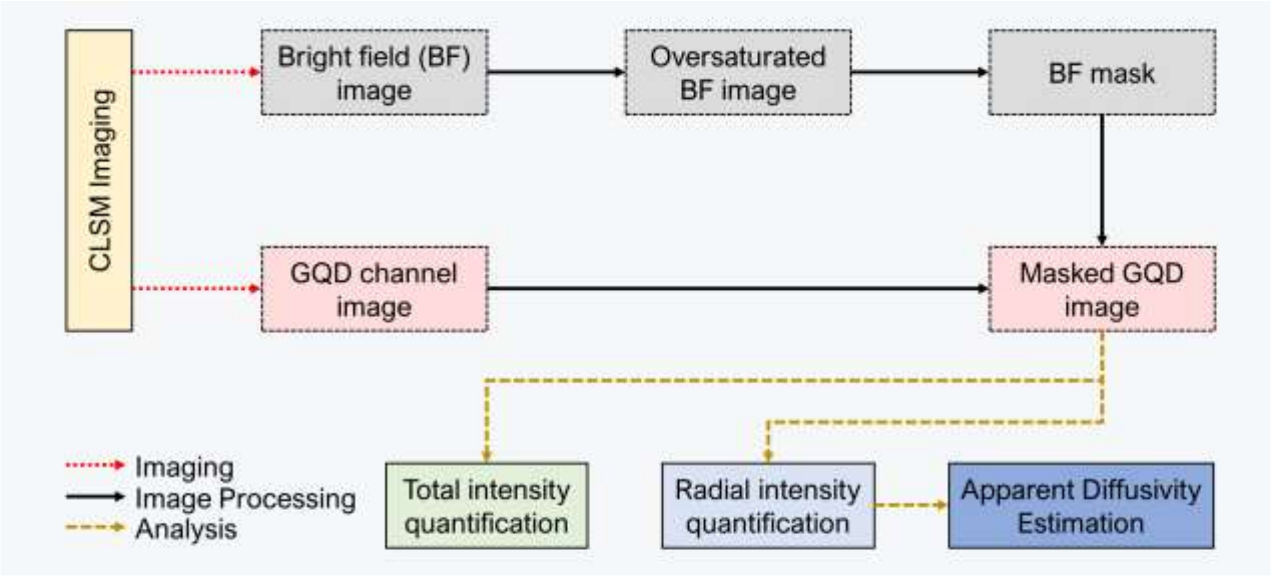


**Supplementary Figure 6.** The workflow for GQD signal collection and quantification from CLSM images.

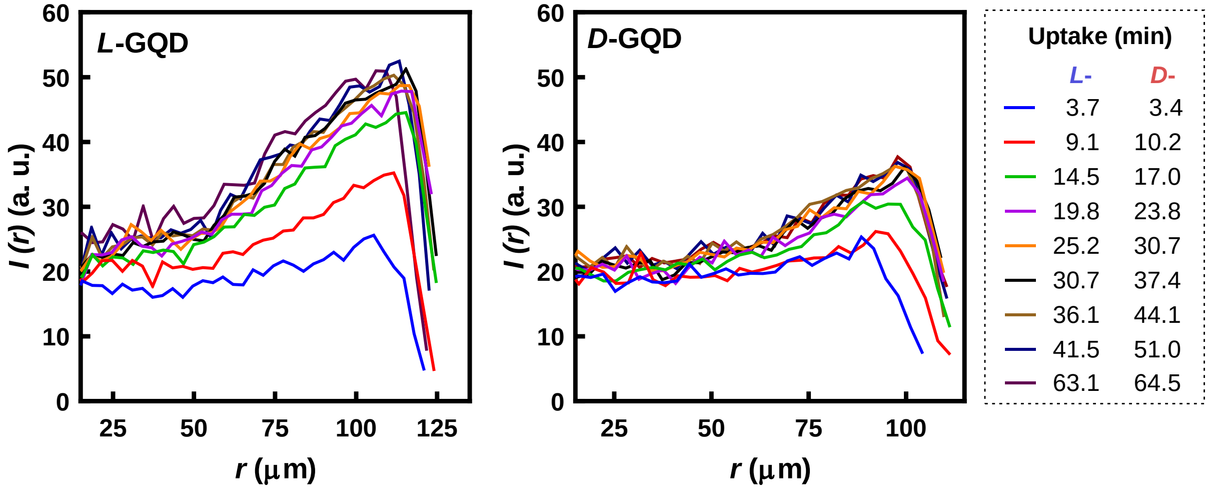


**Supplementary Figure 7.** The average GQD intensities as a function of radius of cellular spheroids under radial coordinates.

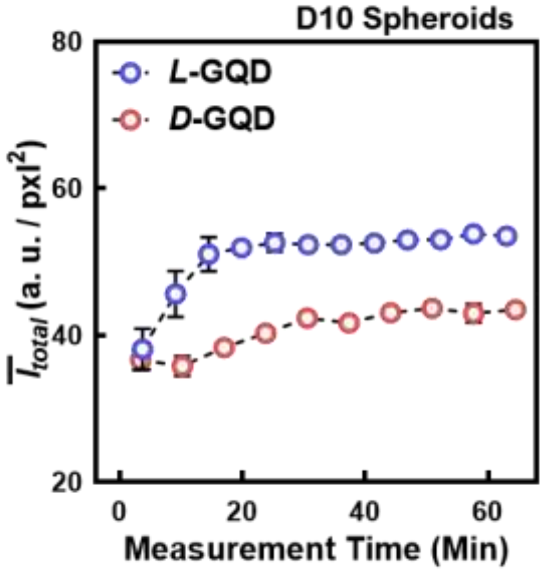


**Supplementary Figure 8.** Integrated GQD intensities increased within the region of cellular spheroids (*e.g.*, 10-day-cultured cellular aggregates) as a function of measurement time (min). Blue curve shows the result from *L*-GQD-treated spheroids, while the red curve shows the result from *D*-GQD-treated spheroids (*N*=3).

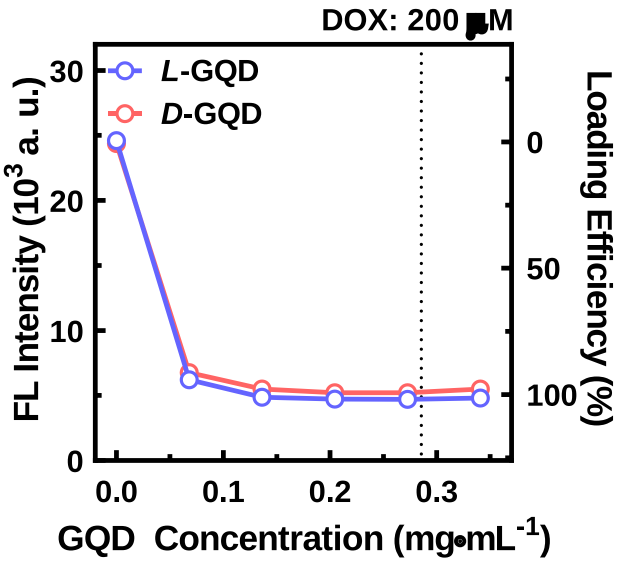


**Supplementary Figure 9.** Fluorescence-based DOX loading efficiency test using *L/D*-GQDs. The loading efficiency was determined by measuring DOX fluorescence at different *L/D*-GQD concentrations, which induced quenching of DOX. Fluorescence intensity was measured at 600 nm emission under 480 nm excitation.

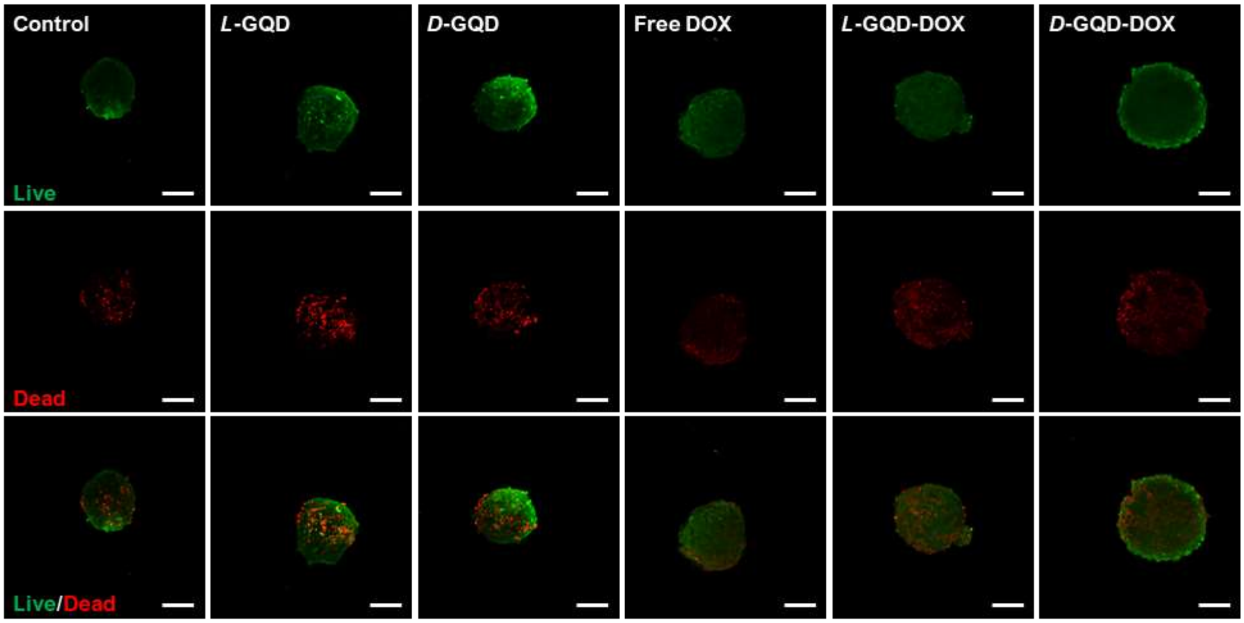


**Supplementary Figure 10.** The confocal fluorescent images of DOX-loaded *L/D*-GQD treatment to 3D cellular spheroid stained with Live/Dead assay (Green: live cell indicator, Red: dead cell indicator, Scale bar: 200 µm).

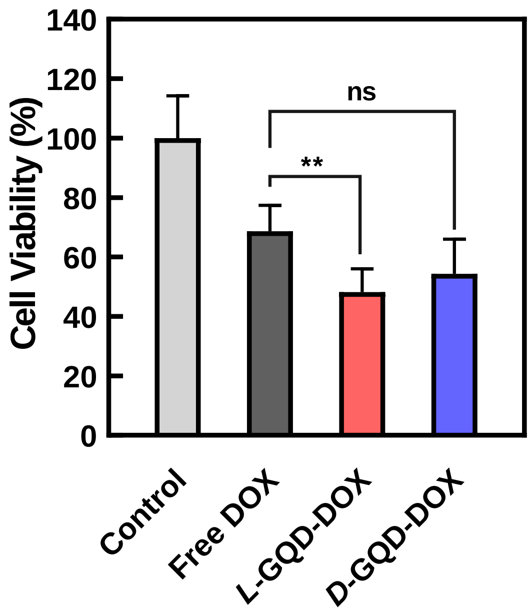


**Supplementary Figure 11.** The result of luminescence-based cell viability in cellular spheroids upon DOX and *L*/*D*-GQD-DOX. The spheroids were exposed to free DOX, *L*-GQD-DOX, or *D*-GQD-DOX solutions (*e.g.*, Prepared as GQD:DOX=0.5 mg·mL$^{-1}$:350 μM; Diluted with fresh cell culture media for 120 μM DOX) for 6 h, followed by the 24-h incubation at 37$^{o}$C (*N*=6).

**Supplementary Table 1.** Wavenumbers of functional groups embedded in *L/D*-GQDs.

| Functional Group | -OH | -C=O | -C=C | C-N | C-O |
|---|---|---|---|---|---|
| Wavenumber ($cm^{-1}$) | 3400<br>1409<br>1350 | 1712 | 1602 | 1250 | 1119 |
